# Negative affective traits moderate tDCS effects on memory

**DOI:** 10.1101/2025.08.15.670509

**Authors:** Marko Živanović, Jovana Bjekić, Carlo Miniussi, Walter Paulus, Saša R. Filipović

## Abstract

**Background:** The cognitive effects of transcranial direct current stimulation (tDCS) show substantial inter-individual variability, emphasizing the need to identify moderators of responsiveness, with negative affectivity as a potential factor influencing both cognitive performance and tDCS outcomes. Here we examine if and how negative affectivity (depressiveness, anxiety, and stress level), conceptualized as both transient states and stable traits, moderates the effects of tDCS on working memory (WM) and associative memory (AM).

**Methods:** We pooled data from six sham-controlled experiments involving 144 healthy young adults (351 tDCS sessions) using within-subject crossover designs. Participants completed WM and AM tasks following active anodal tDCS or sham, as well as Depression, Anxiety, and Stress Scale (DASS-21) before each session.

**Results:** Trait-level, but not state-level, negative affectivity moderated WM. Individuals with higher levels of depression, anxiety, or stress demonstrated greater tDCS-induced WM gains. tDCS-induced AM benefits were consistent and unaffected by affective traits or states.

**Conclusion:** Negative affectivity may shape individual susceptibility to tDCS, increasing the potential for stimulation-induced improvement on tasks requiring significant executive control.

## Introduction

Over the past two decades, interest in non-invasive brain stimulation techniques has grown significantly due to their potential for probing and modulating cognitive functions. Among these techniques, low-intensity transcranial direct current stimulation (tDCS) has significantly advanced our knowledge of the neural mechanisms underpinning cognition. The underlying premise of applying tDCS for cognitive neuromodulation is that delivering electrical current through scalp electrodes can influence cortical excitability, promote targeted circuit activity and neuroplasticity, ultimately modulating cognitive performance, either facilitating or attenuating it[1,2]. The high usability and favorable safety profile[3], coupled with the ability of tDCS to enhance cognitive functions, has led to its investigation in both healthy and clinical populations, with studies reporting improvements in attention[4,5], executive functions[6], learning, and memory[7,8] among other cognitive domains. However, evidence for tDCS efficacy in cognitive enhancement, especially in healthy young adults, remains controversial, with some early meta-analyses showing little to no reliable evidence of behavioral and neuropsychological effects (e.g.,[9]), while other subsequent works depicted a more granulated picture of functions that are more or less susceptible to tDCS effects as well as conditions that either promote or impede neuromodulatory effects of tDCS on cognition in healthy adults[10–12].

Among the cognitive domains, associative memory (AM) and working memory (WM) have been of particular interest due to their fundamental roles in learning, problem-solving, decision-making, and overall daily functioning[13,14]. WM supports the temporary storage and manipulation of information[15], while AM refers to the ability to link previously unrelated pieces of information into complex representations[16]. Since the first study on tDCS effects on WM in healthy adults[17], more than a hundred studies have assessed these effects. One high-quality meta-analysis found overall weak but significant effects of tDCS on WM, attributing inconsistent and seemingly conflicting findings to various factors, including publication bias and low statistical power[18]. More recent meta-analyses on tDCS and WM emphasize the relevance of more specific methodological, neuroanatomical, and measurement-related factors, such as electrode placement, tDCS protocol (online/offline, duration, intensity), electric field distribution, and the type of outcome measures employed, in shaping cognitive outcomes[6,10,19]. Similarly, initial tDCS studies aiming to facilitate AM reported positive effects[20,21], but these were soon followed by experiments that failed to conceptually replicate such findings and reported no significant effects[22,23]. The first meta-analysis of tDCS impact on episodic memory, including AM in healthy adults[24], reported small to negligible overall benefits, highlighting that these outcomes might be largely obscured by substantial methodological heterogeneity. The most recent systematic review specifically focusing on AM[25] found that targeted brain area (i.e., parietal *vs* frontal) and type of outcome measure (recognition *vs* cued-recall) may be the most important differentiators between null and positive results presented in the literature. Overall, positive results (i.e.,improved performance) are more likely when the stimulation is applied over the parietal cortex, and the effects are assessed in cued recall paradigms.

The effects of tDCS show high variability, meaning that they can vary depending on the specific stimulation parameters such as duration, intensity, electrode montage[26], characteristics of the study participants, including neuroanatomical[27], neurochemical[28], genetic[29], and sex differences[30], but also factors such as, the neural excitability state[31], or properties of the outcome measures relative to the study population, such as task difficulty [32,33]. Furthermore, the tDCS effects have been argued to be dependent on experimental conditions, such as successfulness of blinding [34] and prior expectations of tDCS efficacy[35]. Overall, these sources of variability can be split into participants’ characteristics, variable state-based factors, and experimental/contextual factors[36].

A recent umbrella review of meta-analyses examining the impact of prefrontal tDCS across various cognitive domains found that only about a quarter of comparisons demonstrated significant effects, with most evidence classified as low or very low quality[37]. These findings suggest the need to identify potential moderators – variables that could account for the significant variability in observed cognitive outcomes beyond neuroanatomical and experimental factors. Recently, increasing attention has been directed toward psychological characteristics of participants, both their transient state at the time of stimulation (e.g., psychological state) and more stable individual differences (e.g., psychological traits).

In this context, one particularly relevant individual characteristic is negative affectivity, which is characterized by a predisposition to experience negative emotions such as anxiety, stress, and sadness[38]. Negative affectivity can be considered both as a state (momentary emotional experiences) and as a trait (a stable dispositional characteristic), both of which have been linked to cognitive performance[39–42]. From a neuroscience perspective, cognitive and emotional processes are closely interrelated through dynamic interactions of distributed brain regions[43]. Individuals with high negative affectivity exhibit altered functional connectivity and neural responses in key brain regions implicated in cognitive control and memory, such as the prefrontal and parietal cortices[40,44]. Since the prefrontal and parietal cortices are primary targets for neuromodulation in both WM and AM studies, understanding how tDCS effects interact with affective states and traits is crucial.

Given the inconsistent effects of tDCS on cognitive functions, particularly WM and AM, exploring the interaction between tDCS-induced cognitive changes and emotional factors may help in understanding the source(s) of variability in cognitive neuromodulation. A recent meta-analysis revealed that the effects of prefrontal tDCS on executive functions are significantly moderated by mood, further supporting the hypothesis that emotional factors play a key role in neuromodulatory efficacy[45]. Additionally, the concept of affective-state dependency in tDCS has been proposed as a framework[46] for exploring how affective traits and states interact with cortical activity to influence cognitive functions. However, it remains unclear if these effects are due to within-person variations in emotional *state* or whether the negative emotionality represents a relevant, relatively stable *trait* that influences susceptibility to neuromodulatory effects. Our study represents a direct testing of this framework prediction that affective profiles act as internal contextual variables capable of shaping tDCS outcomes through cortical gating mechanisms.

This paper aims to explore the role of negative affectivity in shaping the cognitive effects induced by tDCS. Specifically, we present a comprehensive secondary analysis of multiple experiments examining the effects of tDCS on WM and AM, focusing on how emotional state and trait-like negative affectivity influence these effects. To this end, we pooled together data from six memory experiments[33,47,48] to analyze how trait and state-level of negative affectivity (depression, anxiety, and stress) moderate the effects of tDCS on WM and AM performance.

## Method

### Study Design

The data from six experiments encompassing a total of 351 tDCS sessions (207 anodal active tDCS and 144 sham tDCS) were reanalyzed. All six experiments employed a sham-controlled, within-subject design with counterbalanced order of stimulation conditions and used parallel forms of tasks for repeated assessments. In all experiments, participants underwent each session separately, with a minimum of one week between sessions. Table 1 presents an overview of the main methodological aspects of the studies.

**Table 1.** Description of studies.

*Description of studies*
|  | Study | Sample | Stimulation protocol | Anode placement | Current intensity | Task |
| --- | --- | --- | --- | --- | --- | --- |
| Exp. 1 | Bjekić et al. (2019) | 20<br>($M_{age} = 26.40$, $SD_{age} = 3.71$, 11 females) | offline | P3 | 1.5mA | Face-word (AM) |
| Exp. 2 | Bjekić et al. (2019) | 21<br>($M_{age} = 24.15$, $SD_{age} = 2.74$, 12 females) | offline | P4 | 1.5mA | Object-location (AM) |
| Exp. 3 | Živanović et al. (2021) | 21<br>($M_{age} = 26.76$, $SD_{age} = 4.83$, 12 females) | offline | F3<br>P3 | 1.8mA | Verbal 3-back (WM) |
| Exp. 4 | Živanović et al. (2021) | 21<br>($M_{age} = 26.43$, $SD_{age} = 4.78$, 12 females) | offline | F4<br>P4 | 1.8mA | Spatial 3-back (WM) |
| Exp. 5 | Živanović et al. (2021) | 21<br>($M_{age} = 24.90$, $SD_{age} = 2.49$, 11 females) | online | F3<br>P3 | 1.5mA | Verbal 3-back (WM)<br>Spatial 3-back (WM) |
| Exp. 6 | Živanović et al. (2022) | 40<br>($M_{age} = 25.15$, $SD_{age} = 3.65$, 25 females) | online | P3 | 1.5mA | Short-term associative memory (AM) |

*Note.* The return electrode (i.e., cathode) in all experiments was placed on the contralateral cheek.

Experiments 1, 2, and 6 consisted of two sessions – one active tDCS and one sham, while Experiments 3–5 involved participants going through three experimental sessions – two active tDCS and one sham. In Experiments 1 and 6, tDCS was delivered over the left posterior parietal cortex (PPC), while in Experiment 2, the tDCS was applied over the right PPC. In Experiments 3 and 5, the tDCS was delivered either over the left dorsolateral prefrontal cortex (DLPFC) or left PPC, while in Experiment 4, tDCS was applied either over the right DLPFC or right PPC. So, at the beginning of the session, in Experiments 1, 2, and 6, two electrodes were placed on the participant’s head, whereas in Experiments 3–5, three electrodes were positioned on the participant’s head.

### Participants

Six memory experiments included a total of 144 young, healthy participants (age range: 21–35 years; 57.6% females). Experiment 1 included 20 participants, Experiments 3–5 enrolled 21 participants each, while Experiment 6 included 40 participants. All participants met the inclusion criteria for tDCS[3,49], were right-handed, and had normal or corrected-to-normal vision. Each study was approved by the Institutional Ethics Board, and all procedures adhered to the Declaration of Helsinki.

### tDCS parameters

In all experiments, the electrode of interest delivered anodal stimulation, either in the active or sham condition. In Experiments 1, 2, 5, and 6, a constant current intensity of 1.5mA was used, while in Experiments 3 and 4, the current intensity was 1.8 mA. In Experiments 1 to 5, square 5 x 5 cm electrodes (25 cm^2^) were used, while in Experiment 6, round electrodes also of 25 cm^2^ (_≈_ 5.6 cm diameter) were used. In all experiments, the rubber electrodes were inserted into saline-soaked sponge pockets before positioning on the scalp.

In Experiments 1 and 6, the active or sham tDCS electrode was placed over the left PPC (P3), while in Experiment 2 it was placed over the right PPC (P4). In Experiments 3–5, the electrode location was randomized across sessions and targeted either the DLPFC (left i.e.,F3 in Experiments 3 and 5; right, i.e.,F4 in Experiment 4) or the PPC (left i.e. P3 in Experiments 3 and 5; right, i.e.,P4 in Experiment 4). In all experiments, the return (i.e.,cathodal) electrode was placed on the contralateral cheek. Therefore, WM experiments applied tDCS over both DLPFC and PPC, whereas AM experiments used only the PPC montage. Figure 1 illustrates the electrode placements and e-field simulations. The estimated e-field strength in the target regions was relatively consistent across montages (*M*=0.290, range 0.204–0.320;see Supplement 1).

**Figure 1.**
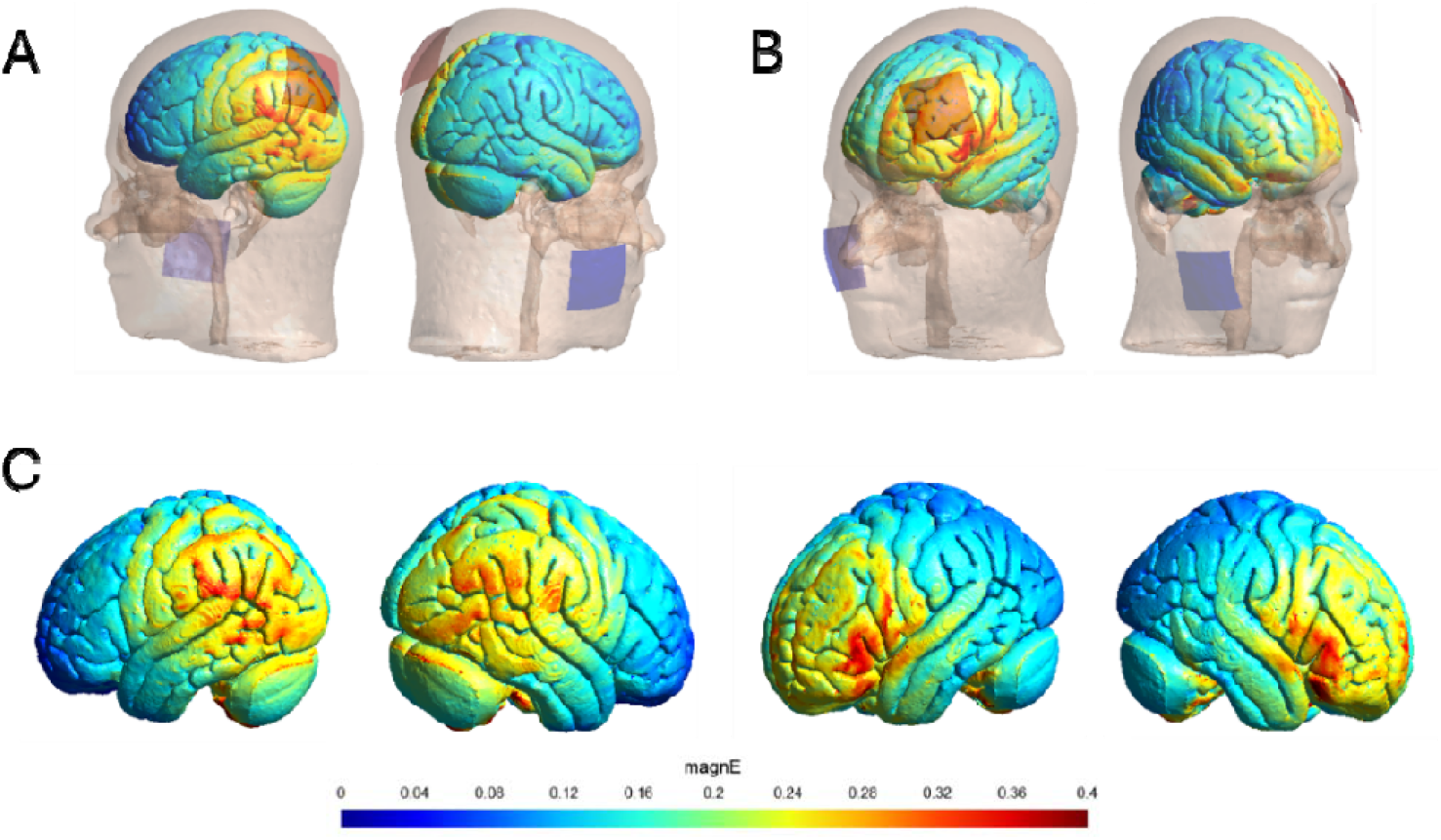
Representation of electrode placement and estimated induced e-fields across montages. (A) PPC montage: anode over left PPC (P3) and cathode over the contralateral cheek. (B) DLPFC montage: anode over left DLPFC (F3) and cathode over the contralateral cheek. (C) Estimated e-field distributions for left PPC, right PPC, left DLPFC, and right DLPFC.

In all six experiments, the active tDCS condition was delivered for 20 min, using a 30s ramp-up and 30s ramp-down period. In sham conditions, the electrode montage was kept the same to the active tDCS condition, and the current was only briefly delivered at the beginning and at the end of the 20min in a ramp-up/down manner (i.e., over the first and last 60s).

### Memory Assessment

In Experiment 1, AM was assessed using a face-word memory task[47]. The task consisted of two learning blocks, each followed by a test block. During the learning block, participants were successively presented with 20 face-word pairs and instructed to memorize as many as possible. In the test block, the pictures of the same 20 faces (“old faces”) were intermixed with 30 new faces and successively presented in quasi-random order, and the participants were asked to recognize the faces from the previous block. For each face recognized as “old”, participants were asked to write in the word that a given face had been previously paired with. After the first test block, the participants had another opportunity to learn the same face-word pairs in the second learning block, followed by another face-cued word recall test block. The percentage of correctly recalled face-word associations in both test blocks served as the outcome measure for this task.

In Experiment 2, AM was assessed using the object-location associative memory task[47]. The task consisted of two learning blocks, each followed by a test block. In the learning block, 15 colored images of everyday objects (e.g., furniture) and living things (e.g., animals) were sequentially, quasi-randomly presented in a 4x4 grid. Participants were instructed to memorize the locations of each stimulus in the grid. In the test block, the same images were shown above the grid, and participants needed to recall the location in which each stimulus had been previously presented. As in the previous task, in the second learning block, participants had another opportunity to remember object-location associations, which was, again, followed by a test block. The percentage of correctly recalled locations in both test blocks was used as the outcome measure.

In Experiments 3–5, WM was assessed using verbal and spatial 3-back tasks[48]. Both tasks had a similar structure, consisting of one training and five test blocks. In the verbal 3-back task, ten Latin alphabet letters were sequentially presented on the screen. Participants were instructed to respond by keypress when presented with the same letter as in the three steps before. In the spatial 3-back task, participants were shown a black square that successively changes its position in the 3x3 white grid. They were instructed to press a key when the black square was in the same position as in the three trials before. In both tasks, 25% of all trials required participants to react. Outcome measures in verbal and spatial 3-back tasks were calculated as the percentage of hits (i.e., correctly identified targets).

In Experiment 6, AM was assessed using the short-term AM task[33]. The stimuli in this task consisted of single digits (0-9) printed in white, sequentially presented in fixed pre-randomized order on backgrounds of different colors (blue, green, red, yellow, pink, or grey). Digit-color associations were presented to participants in sequences of different lengths, varying between 3 and 5 associations, and the participants were instructed to remember each digit-color association. At the end of each sequence, participants were presented with one of the previously seen color cards and were instructed to recall the digit that was presented on a given color. Since the target digit could appear at each position in a sequence, its frequency was balanced across sequences. The measure of AM was calculated as the percentage of correctly recalled digits.

Across all six experiments, parallel forms of the memory tasks were counterbalanced across sessions and experimental conditions.

### Negative affectivity questionnaire

The Depression Anxiety Stress Scales (DASS-21) is a 21-item self-report scale designed to assess depression, anxiety, and stress, with 7 items dedicated to each domain [50]. Participants rate each item using a 4-point scale indicating how much each statement applied to their mood (0 – did not apply to me at all, 3 – applied to me very much or most of the time) over the past week. For each domain and full-scale DASS, scores were calculated by averaging participants’ responses on corresponding items (ranging from 0 to 3).

### Procedure

In Experiments 1–4, behavioral effects were assessed in the so-called *offline* protocol, i.e., immediately after the stimulation. In Experiments 5 and 6, the *online* protocol was employed, i.e., the effects were assessed during tDCS. In each experiment, DASS-21 was administered at the beginning of each tDCS session, i.e., before active tDCS or sham conditions. Before and after tDCS application, a standard questionnaire [51] was administered to monitor for potential side effects and adverse events. No serious adverse events were recorded, confirming good safety and tolerability of the procedure; tDCS induced only mild discomfort and mainly typical sensory effects (e.g., tingling) (for details see [33,47,48]). In each experiment, end-of-the-study sham guessing was adopted to evaluate the successfulness of blinding. Blinding was successful across all experiments [33,47,48], and a separate study, which analyzed Experiments 1–4, showed that successful sham guessing did not moderate tDCS effects[52].

Each session lasted approximately 45min in Experiments 1–4, and about 30min in Experiments 5 and 6. Accounting for electrode setup and the completion of all questionnaires, the total duration of each session was around 1h.

### Data analysis

Data from six memory experiments were aggregated into two datasets–one for WM (*N*=63 participants; 378 observations) and one for AM (*N*=81 participants; 163 observations); providing a stronger basis for reliable statistical analyses. As in the initial publications, WM data were analyzed as hit rates centered by session order [48], while AM data were analyzed as the proportion of successful cued recall. All analyses were conducted using IBM SPSS Statistics 21, JASP0.95, and R4.5.1. The dataset is openly available on the Open Science Framework (OSF) at https://osf.io/g3my7/?view_only=ad99f16b160f4ba9a9480ad0f3e8dddd.

To assess the *differences* between active tDCS and sham conditions in negative affectivity (depression, anxiety, stress, DASS-21 total scores) across experiments, a series of repeated-measures ANOVAs was conducted. To quantify cross-condition *consistency* in negative affectivity, we calculated test-retest correlations (Pearson *r*) and intraclass correlation coefficients (*ICC*, two-way random effect method, absolute agreement type, average measures) between the corresponding active tDCS and sham conditions. *Variance decomposition* of negative affectivity was performed to distinguish trait-like stability from state-like fluctuations. To decompose the reliable variance, we used test-retest reliability (*r*_*tt*_) reflecting stable reliable variance and internal consistency measures of negative affectivity scales reflecting total reliable variance (Cronbach’s α). Specifically, we computed the percentages of reliable variance in negative affectivity attributable to relatively stable trait-like individual differences using the formula *r*_*tt*_/α_max_*100, as a conservative proxy for trait-level (between-person) variability. The variance attributable to state-level (within-person) variability was calculated as (1–*r*_*tt*_/α_max_)*100.

Measures of relatively stable, trait-like individual differences for each domain of negative affectivity were calculated as the average scores of all conditions, while state-level fluctuations were computed as deviations from the mean i.e., trait-level.

To examine the relationship between state fluctuations in negative emotions and tDCS-induced changes in WM and AM performance, we calculated Pearson’s correlation coefficients between person-mean centered scores of each negative affectivity domain and tDCS-induced gains in memory performance. To examine the relationship between stable individual differences in negative emotions and memory performance, we calculated Pearson’s correlation coefficients between each average negative affectivity domain (across all conditions) and tDCS-induced gains in memory performance (active memory performance tDCS – sham memory performance). Additionally, to determine if measures of negative affectivity and memory performance are related regardless of stimulation condition, we correlated average negative affectivity scores with average memory performance across conditions.

The core inferential analyses involved a series of linear mixed models (LMM) testing if state-level fluctuations or trait-level negative affectivity influence and moderate initially observed tDCS effects on WM and AM. In all models, CONDITION (active tDCS *vs* sham) was used as a fixed factor while participants’ ID was used as a random grouping factor (e.g., *WM_score∼stimulation_condition+(1*|*ID)*). Separate models were specified for each domain of negative affectivity, with state-level and trait-level moderators entered simultaniously, (e.g., *WM_score∼stimulation_condition*(anxiety_state+anxiety_trait)+(1*|*ID)*).

Continuous moderators were probed at three levels: low (1*SD* below the mean), mid (moderator’s mean), and high (1*SD* above the mean). All post hoc comparisons were conducted using Satterthwaite’s method for degrees of freedom estimation and were corrected for multiple comparisons using the Holm procedure. Degrees of freedom were estimated by Satterthwaite’s approximation for fixed effects, and by Kenward–Roger method for post-hoc comparisons, with multiple comparisons control using Holm procedure applied within each affectivity domain.

Exploratory, we tested more comprehensive multilevel models, with factors TASK, LATERALITY, and PROTOCOL type (Supplement 6–7). Furthermore, models with random intercepts for study and random slopes for condition at the participant level were tested but turned out to be over-specified for the data (singular fits). The absence of effects of other potential confounding factors, such as session order, between-session learning, subjective experience, and stimulation-related sensations, as well as blinding success, has been reported in previous publications [33,47,48].

## Results

Descriptive statistics for three DASS-21 subscales and overall scores (Supplement 2) showed that participants, on average, showed relatively low negative affectivity, consistent with expectations for healthy young adult samples. WM and AM measures showed the expected range and well-calibrated difficulty across tasks and conditions (Supplement 3).

### Stability of negative affectivity

Across experiments, no significant differences in DASS-21 subscales between active tDCS condition(s) and sham were found, with the only exception of slightly elevated depression scores before active tDCS in AM experiments. Still, only moderate stability across conditions was observed, as most *ICCs* were below .80 and test-retest correlations ranged between .43 and .71 (Table 2).

**Table 2.**
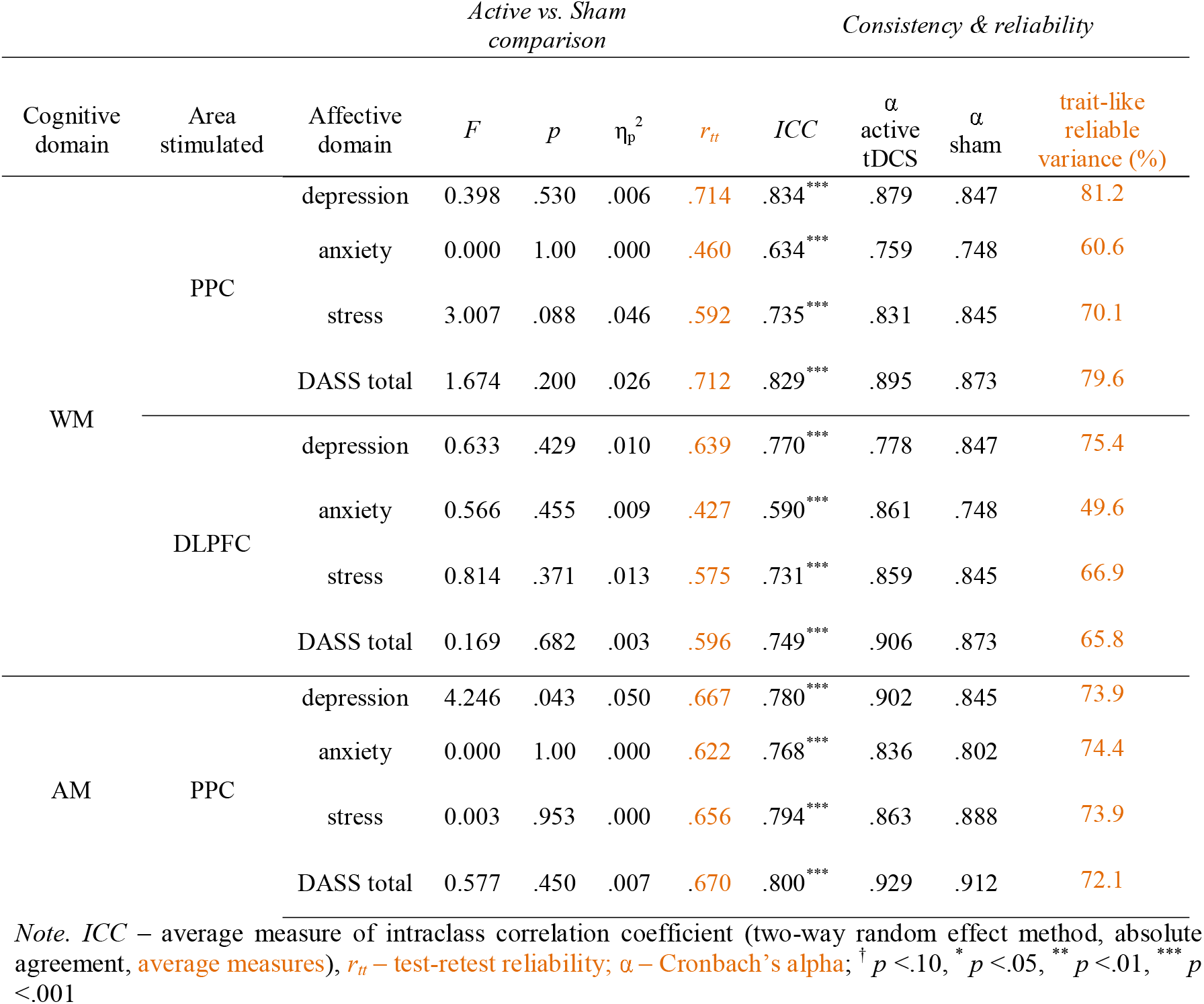
Differences and consistency in negative affectivity domains before active tDCS and sham stimulation, with reliability indices (test-retest, Cronbach α), and % of variance attributable to stable individual differences, across cognitive domains and areas stimulated

The highest consistency in negative affectivity between active tDCS and sham conditions was observed for depression, followed by stress, with the lowest consistency found for anxiety. On average, ICCs were .795 for depression, .753 for stress, and .664 for anxiety. These discrepancies in consistency could only partly be attributed to scales’measurement error, as median reliability was highest for stress (*Mdn* α=.859), and depression (*Mdn* α=.847), followed by anxiety (*Mdn* α=.802). After correcting for measurement error, 76.9% of the true variance in depression, 70.3% in stress, and 61.5% in anxiety reflected stable interindividual differences, leaving the remainder of true variance to the state-level variations (Supplement 4).

### Relations between negative affectivity and cognitive performance

At the level of all observations (regardless of stimulation condition), domains of negative affectivity showed significant negative correlations with WM performance across all tDCS conditions (Table 3). However, when *state-level* variations were considered, regardless of the area stimulated, no significant correlations were observed between tDCS-induced WM-gains and state-level variations in affectivity. In contrast, on a *trait-level*, anxiety, stress, and DASS total score were all significantly and positively related to tDCS-induced WM-gains following stimulation of PPC. A similar pattern was observed following DLPFC stimulation, with higher tDCS induced WM gain observed in higher trait stress levels and a higher DASS total score, while the same trend was observed for anxiety. The findings indicate that almost all shared variance between WM performance and negative affectivity can be attributed to relatively stable factors. In addition, the average WM performance (across all stimulation conditions) was positively related to anxiety and DASS total score, and on a trend level, with depression as well. On the other hand, AM performance was not related to negative affectivity on both the state and trait levels.

**Table 3.**
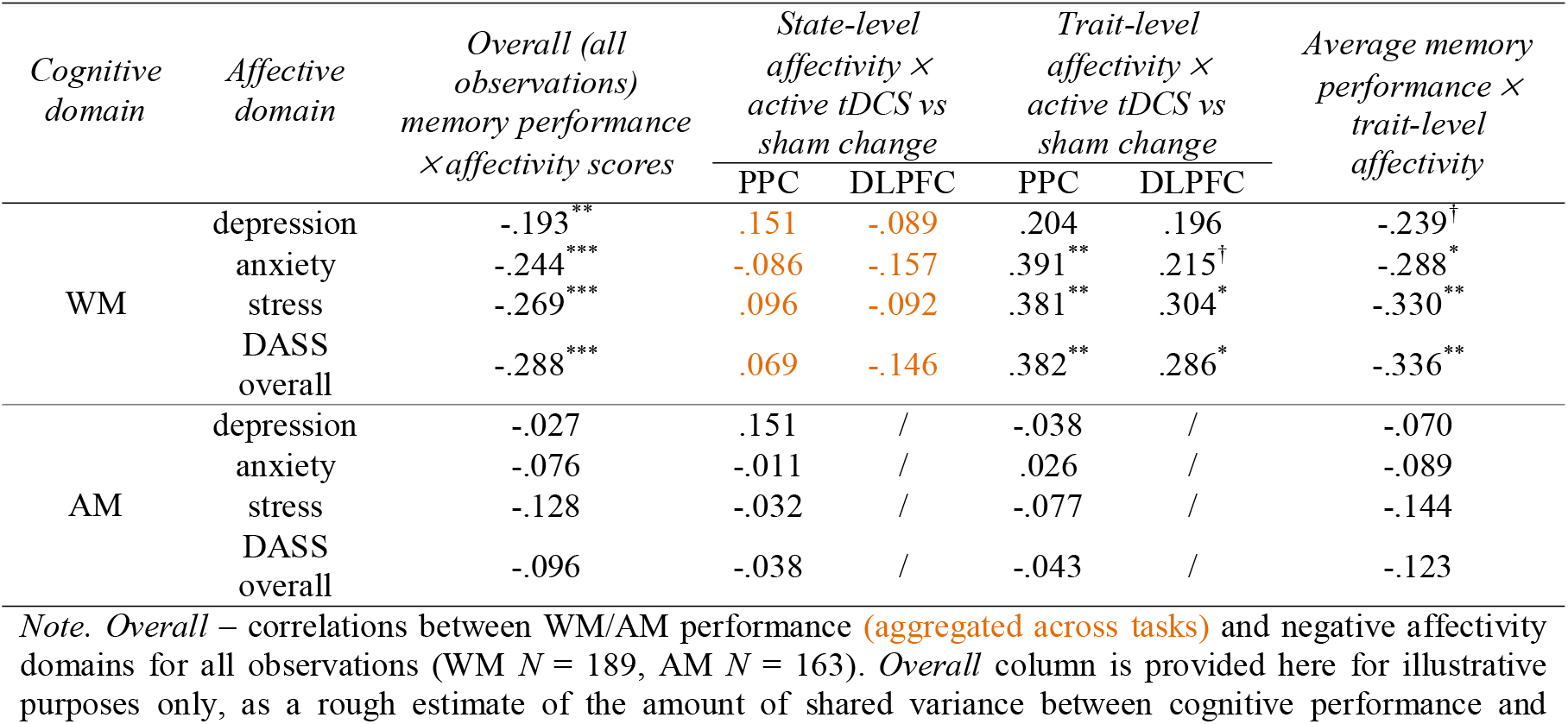
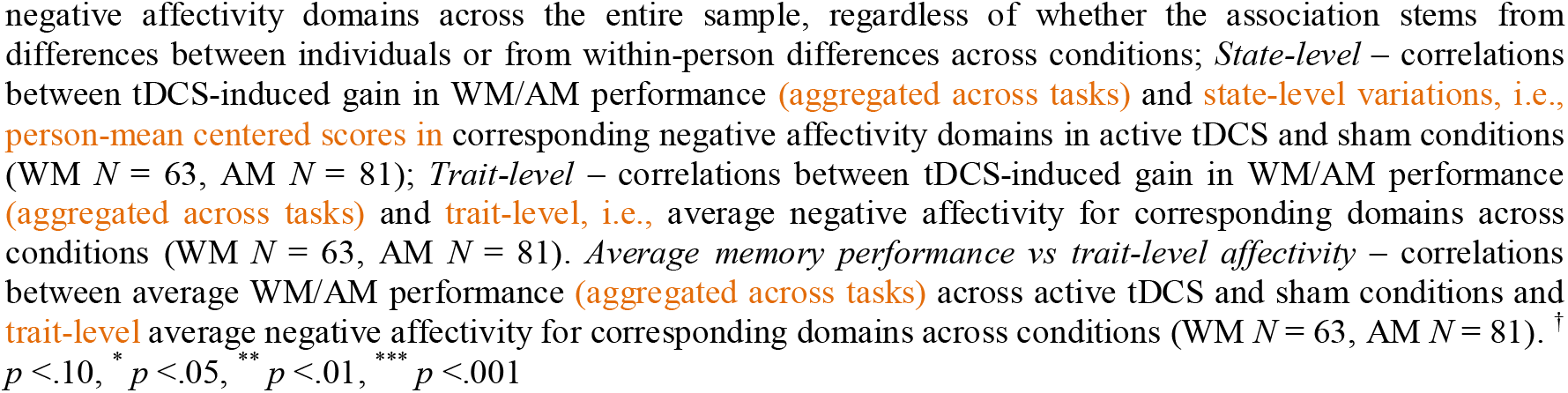
Pearson correlations between negative affectivity and WM and AM cognitive performance on state- and trait-levels.

### Negative affectivity as a moderator of tDCS effects

#### Working memory

Based on the combined dataset from the three WM experiments, results of the LMMs revealed an overall significant main effect of CONDITION on WM performance [*F*_(1,314)_=5.216, *p*=.023, η_p_^2^=.02]. Initial tDCS effects on WM performance were no longer significant once state- and trait-level negative affectivity were included as moderators: depression [*F*_(1,310.94)_=0.882, *p*=.348], anxiety [*F*_(1,311.19)_=0.003, *p*=.960] stress [*F*_(1,311.58)_=1.285, *p*=.258]and DASS total [*F*_(1,311.33)_=0.708, *p*=.401]. Significant main effects emerged only for state-level anxiety [*F*_(1,323.95)_=4.229, *p*=.041, η_p_^2^=.01], whereas trait-level effects were observed for all negative affectivity measures including depression [*F*_(1,62.20)_=4.799, *p*=.032, η_p_^2^=.07],anxiety [*F*_(1,62.84)_=7.272, *p*=.009, η_p_^2^=.10], stress [*F*(1,62.58)=9.501, *p*=.003, η_p_^2^=.13], and DASS total [*F*_(1,62.56)_=9.893, *p*=.003, η_p_^2^=.14]. Noteworthy, the initial tDCS effect of WM remained significant in the model that included only state-level negative affectivity (Supplement 5).

For state-level, no significant STIMULATION X NEGATIVE EMOTION interactions were observed for anxiety *F*_(1,355.49)_=0.182, *p*=.670, stress *F*_(1,369.76)_=0.909, *p*=.341, DASS total score *F*_(1,367.32)_=2.010, *p*=.157], while state-level depression showed marginal effect *F*_(1,367.84)_=3.975, *p* =.047, η_p_^2^=.01.

However, for trait-level significant CONDITION X NEGATIVE EMOTION interactions were observed for anxiety [*F*_(1,311.64)_=9.422, *p*=.002, η_p_^2^=.03], stress [*F*_(1,311.25)_=10.608, *p*=.001, η_p_^2^=.03], and DASS total [*F*_(1,311.18)_=10.085, *p*=.002, η_p_^2^=.03], while a marginal effect was found for CONDITION X depression interaction [*F*_(1,310.87)_=3.769, *p*=.053, η_p_^2^=.01]. In other words, trait-level negative affectivity moderated the effects of active tDCS on WM performance. For each negative emotion, significant Holm corrected tDCS effects were observed for mid and higher, but not lower levels of trait [depression: mid *t*_(311)_=2.423, *p*=.032, higher *t*_(311)_=3.082, *p*=.007; anxiety: mid *t*_(311)_=2.422, *p*=.032, higher *t*_(311)_=3.869, *p*<.001; stress: mid *t*_(311)_=2.337, *p*=.040, higher *t*_(311)_=3.955, *p*<.001; DASS total: mid *t*_(311)_=2.320, *p*=.042, higher *t*_(311)_=3.886, *p*<.001], indicating that tDCS attenuates the negative association between WM performance and negative affectivity (Figure 2).

**Figure 2.**
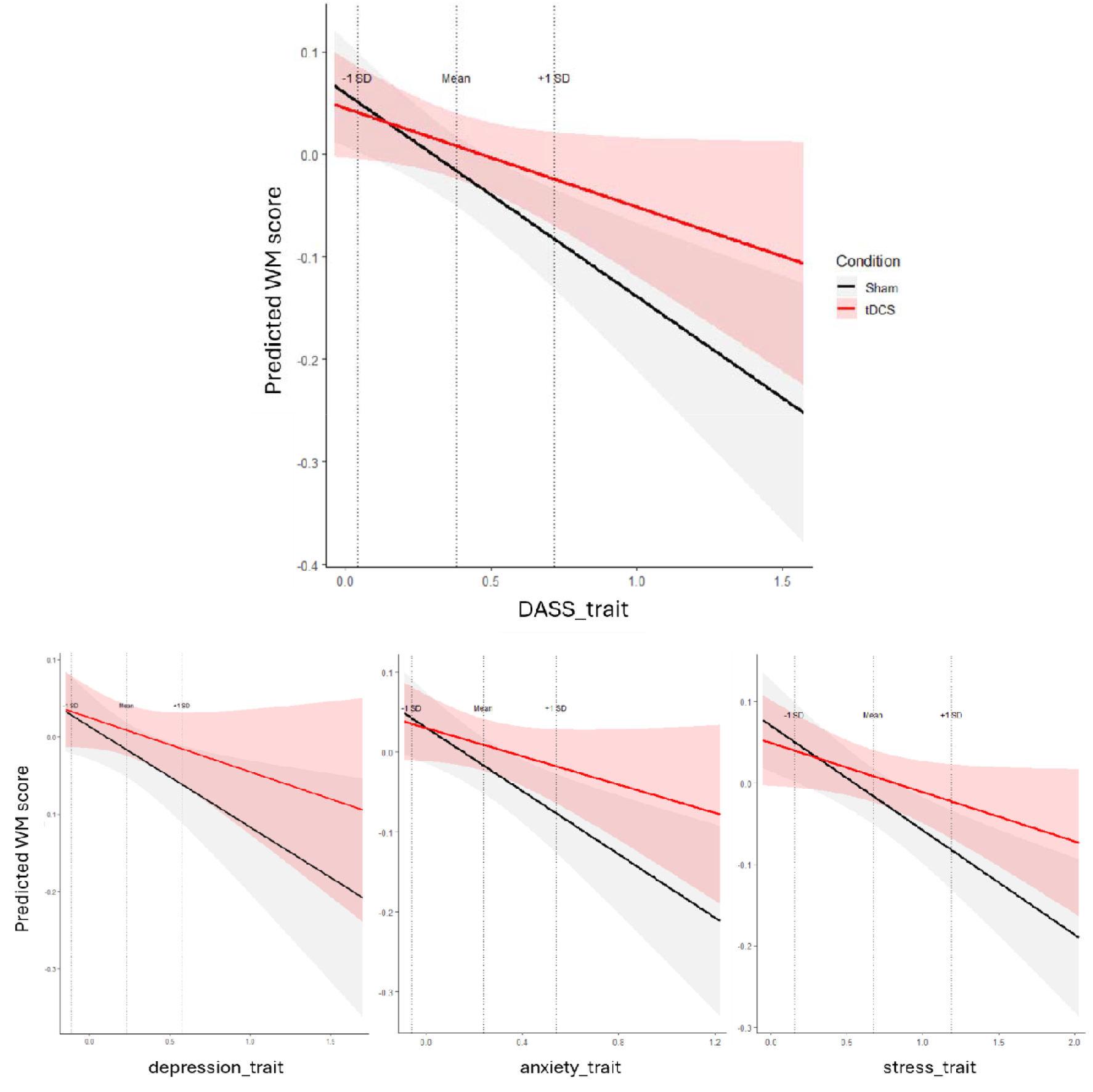
Moderation of tDCS effects on WM by trait negative affectivity. Predicted WM scores (y-axes) are plotted as a function of DASS total score (x-axes) (top panel) and the three subscales: depression, anxiety, and stress (bottom panels). Black lines represent sham and red lines represent tDCS, with shaded areas indicating 95% confidence intervals. Vertical dashed lines denote trait levels at −1 SD, the mean, and +1 SD.

Effect sizes indicated small but consistent benefits at mid-levels (depression: *d*=0.27, 95%CI [0.05,0.48]; anxiety: *d*=0.27, 95%CI [0.05,0.48]; stress: *d*=0.26, 95%CI [0.04,0.48]; DASS total: *d*=0.25, 95%CI [0.04,0.47]) and moderate-to-large effects at higher trait levels (depression: *d*=0.48, 95%CI [0.17,0.78]; anxiety: *d*=0.60, 95%CI [0.30,0.91]; stress: *d*=0.62, 95%CI [0.31,0.92]; DASS total: *d*=0.60, 95%CI [0.30,0.91]).

Additional exploratory LLMs, including task, laterality, and protocol (Supplement 6) showed no moderation by these factors and no systematic deviation from the pattern where trait, but not state, negative affectivity moderated tDCS effects on WM.

#### Associative memory

After pooling together data from three AM experiments, a significant overall stimulation effect on AM was observed [*F*_(1,80)_=15.817, *p*<.001, η_p_^2^=.17]. Initial tDCS effects on AM performance remained significant after introducing state- and trait-level negative affectivity to the model: depression [*F*_(1,78)_=11.291, *p*=.001,η_p_^2^=.13], anxiety [*F*_(1,78)_=8.987,*p*=.004,η_p_^2^=.10], stress [*F*_(1,78)_=8.806, *p*=.004,η_p_^2^=.10], as well as DASS total [*F*_(1,78)_=8.313,*p*=.005, η_p_^2^=.10]. In neither case, significant main effect of state-level negative affectivity domains was observed [depression state *F*_(1,78)_=2.423, *p*=.124, trait *F*_(1,78)_=0.357, *p*=.552; anxiety state *F*_(1,78)_=0.006, *p*=.937, trait *F*_(1,78)_=0.669, *p*=.416; stress state *F*_(1,78)_=0.089, *p*=.766, trait *F*_(1,78)_=1.593, *p*=.211; DASS total state *F*_(1,78)_=0.161, *p*=.690, trait *F*_(1,78)_=1.411, *p*=.238].

Furthermore, no CONDITION X NEGATIVE EMOTION interactions were observed for state level [depression *F*_(1,78)_=0.002, *p*=.964; anxiety *F*_(1,78)_=0.390, *p*=.534; stress *F*_(1,78)_=0.756, *p*=.387; DASS total *F*_(1,78)_=0.597, *p*=.442], nor trait-level [depression *F*_(1,78)_=0.681, *p*=.412; anxiety *F*_(1,78)_=0.054, *p*=.817; stress *F*_(1,78)_=0.518, *p*=.474; DASS total *F*_(1,78)_=0.203, *p*=.654].

In additional exploratory LLMs including TASK, LATERALITY, and PROTOCOL TYPE (Supplement 7), some multifactor interactions emerged; however, the initial effects of tDCS on AM were robust.

## Discussion

Building on growing evidence of high inter- and intra-individual variability in neuromodulatory tDCS effects, in this study we explored the interaction between cognitive and affective processes as a potential source of variability in memory-related tDCS outcomes. By pooling data from multiple tDCS experiments, we demonstrated that in healthy participants, stable emotional traits, but not transient fluctuations in emotional states, moderate the effects of tDCS on WM. Conversely, the effects of tDCS on AM proved to be robust, remaining unaffected by negative emotional states at both state and trait level.

The study addresses an underexplored area in tDCS research, with findings that both align with and extend previous work suggesting that **e**motional factors may shape the neuromodulatory effects of tDCS (for review see[46]), particularly in tasks that rely heavily on attentional and executive control processes. Recent meta-analytic findings have shown that the effects of prefrontal tDCS are significantly moderated by mood, with individuals with higher depressive symptoms experiencing greater benefits in executive functioning from tDCS application[45]. Moreover, the affective-state dependency framework[46] has been proposed as a promising approach to disentangle the dynamic relationships between emotional state, cortical excitability, and cognitive functions. Still, this framework remains largely unexplored and mostly based on indirect evidence, e.g., studies showing differential tDCS effects on emotionally charged stimuli[53], differential tDCS effects in different experimentally-induced emotional states such as high/low stress[54] or anger[55], and the relationship between emotion-regulation strategies and cognitive tDCS effects[56]. Only a few studies assessed the relationship between individual differences in emotions and cognitive tDCS effects. Namely, Esposito et al [57,58] found that responsiveness to tDCS varies depending on the trait and state anxiety levels, while another study found the effects of trait anxiety[59]. Still, the role of other negative emotions beyond anxiety remains unexplored. It is plausible that trait-level affectivity shapes the baseline configuration and variability of cortical network activations that underlie an individual cognitive response within a homeostatic neural space, while momentary emotional states may act as narrow modulators of activity within this dynamic framework. Furthermore, it remains unclear whether affective modulation of cognitive tDCS effects is driven primarily by within-person fluctuations in emotional state or by more stable, trait-like emotional characteristics.

Our data provide evidence in favor of the latter as tDCS effects on WM are shown to be sensitive to trait-level negative affectivity rather than state-level variations. This supports the hypothesis that stable emotional traits, rather than relatively transient variations in emotional states, might shape cortical susceptibility to tDCS in neurotypical adults. To the best of our knowledge, only one study explored the role of relatively stable traits (e.g., proneness to experiencing negative emotions) in the context of responsiveness to tDCS[60], however, that study’s findings are less robust due to a very small sample size. Consistent with a large body of literature[61,62], our data show that negative emotions exhibit considerable stability in young, healthy participants, with the majority of variance in negative affectivity, on average, being attributable to stable individual differences rather than emotional fluctuations. The finding aligns with trait-based models of negative affectivity[38]. This study, using psychometrics to disaggregate variance sources, adds empirical weight to the idea of trait-based susceptibility to tDCS.

Furthermore, we found that stable individual differences in experiencing negative emotions have a differential effect on WM and AM performance. While the cognitive performance that is heavily reliant on executive control processes, such as WM, was consistently and negatively associated with trait-level negative affectivity, this was not the case for associative binding. In contrast, state-level fluctuations in negative emotions were not consistently related to tDCS-induced changes in cognitive performance, for either WM or AM. Thus, only trait-like dispositions toward experiencing negative emotions emerged as significant moderators of tDCS effects of WM. Specifically, individuals with moderate and higher trait levels of negative affectivity showed greater improvements in WM performance following tDCS. On the other hand, tDCS appeared to be ineffective in modulating WM in individuals with low levels of negative affectivity, at least with the stimulation settings used in these experiments. One possible explanation for these findings could be that trait-level negative affectivity may be associated with different baseline cortical excitability states in prefrontal regions involved in the WM task. Individuals with higher trait anxiety or stress could exhibit a modified inhibitory tone, which might potentially enhance the responsiveness of prefrontal circuits to neuromodulatory input[63]. Although speculative in the absence of direct neural measures, this interpretation is consistent with prior evidence showing that anticipatory anxiety can alter cortico-spinal excitability[64].

Taken together, our results suggest that the effects of tDCS on WM processes are influenced by the degree of interference arising from emotional dysregulation. Specifically, tDCS appears to have greater modulatory potential in individuals with less efficient emotional regulation. This may suggest that higher levels of negative affectivity contribute to suboptimal cognitive performance, thus leaving room for improvement by tDCS. Conversely, people with lower levels of trait negative affectivity demonstrated reduced responsiveness to stimulation, suggesting that already optimal cognitive functioning in emotionally stable individuals, with stable homeostatic regulation, limits tDCS enhancing potential. Broader markers of baseline cognitive performance (e.g.,IQ) and neurophysiological measures (e.g.,EEG) need to be examined alongside affective traits to better understand the boundary conditions of tDCS efficacy. Besides influencing psychological processes at the behavioral level, emotional traits also shape underlying neurophysiological variability[65]. Cortical activity in individuals with high negative affectivity is associated with altered functional connectivity and neural responses in prefrontal and parietal cortices[40,44]. This suggests a potential pathway by which emotional traits may influence baseline cortical excitability, which tDCS interacts with to produce varied outcomes.

Moreover, although not directly investigated in this study, our results draw attention to the possibility that some of the cognitive effects reported in the tDCS literature may be either hindered or enhanced through the concurrent neuromodulation of emotional control/processing systems. In one respect, modulation of key nodes within the central executive network is likely to influence not only cognitive functions per se, but also emotional regulation, and vice versa. However, more importantly, a strong modulatory effect on negative affectivity, especially depression, has already been established for DLPFC stimulation and has been harnessed in treatment applications for depression and other conditions associated with negative affectivity [66,67]. The latter issue may not be obvious in neurotypical individuals, but it can still exert a subtle impact on the results of brain stimulation studies on memory and other cognitive functions.

Interestingly, AM performance was not significantly related to state or trait negative affectivity and demonstrated robust and consistent enhancement following tDCS. The stable effect on AM is not attributable to other potential confounding factors, such as expectancy or correct stimulation-condition guessing[33,47,52] or session order[33,47]. This dissociation between WM and AM possibly stems from the neural and cognitive differences between them. Namely, both WM and emotional regulation rely heavily on the top-down control mechanisms of the central executive network. Cognitive performance in attention/resources-demanding WM tasks is particularly influenced by a person’s ability to suppress affective interference. In contrast, AM depends more on the hippocampal-cortical pathways, with critical contribution of the PPC, making it less vulnerable to emotional interference and less affected by the competition between emotional and cognitive processes for limited neural resources.

Together, these findings underscore the significance of task- and person-specific factors in determining the efficacy of tDCS interventions. While WM performance is susceptible to modulation by trait negative affectivity, improvement in AM performance was robust to state and trait level variability in negative emotions. Moreover, it seems that tDCS, in the present work, has limited potential in modulating attention-dependent cognitive processes in emotionally stable individuals who perform at their optimal level.

While our findings offer valuable insights into the interplay of emotions and cognitive tDCS effects, several limitations should be noted. First, we included only studies with healthy young adult participants who are expected to show relatively restricted variability in emotional state. It remains an open question whether these findings would be replicated in populations where both cognitive performance and emotional states vary more substantially, such as an aging population or clinical groups. Moreover, DASS-21 does not capture state in the narrowest sense (i.e., momentary affect), but rather variation over several days, and as such may be susceptible to retrospective bias. Next, to achieve the equivalence of assessment tools, we used data from the same instrument to extract measures of both trait- and state-like negative affectivity. However, this approach also holds its limitations, since there is a certain amount of shared method variance between the measures. Additionally, measures of negative affectivity were collected at two or three time points within a relatively short interval of several weeks. A greater number of assessment points over a longer period would allow for a more reliable estimation of stable individual differences, reducing the influence of transient state-related fluctuations in this measure. In addition, we did not have data to test potential interactions between DLPFC stimulation, negative affectivity, and AM performance. Finally, all experiments were designed as a single-session protocol, yet, to fully capture the interplay between emotional state and tDCS-related cognitive effects, a longitudinal study with more sophisticated methods, such as ecological momentary assessment, would offer a more nuanced insight into the potential interaction of weekly/daily/momentary emotional fluctuations within each person and tDCS-related cognitive outcomes. Nonetheless, the use of six distinct experiments, unified by affective measures, provides a robust foundation for the present analyses.

## Conclusion

The present results show that negative trait-level affectivity significantly moderates the effects of tDCS on WM performance, whereas variations in negative state-level affectivity do not appear to yield a comparable effect. These results emphasize the importance of considering individual differences in psychological traits when evaluating the results of tDCS experiments. Negative affective traits, such as depression, anxiety, and stress, may affect both cognitive performance and baseline cortical excitability, which in turn might promote tDCS effectiveness. This work aligns with recent evidence suggesting that psychological traits can shape the variability of tDCS effects. Moreover, these findings highlight the potential value of integrating affective assessments into future neuromodulation approaches aimed at optimizing cognitive performance.

## Acknowledgement

The authors would like to thank the researchers who contributed to the collection of the dataset analyzed in this study: Dunja Paunović, Uroš Konstantinović, Katarina Vulić, Marija Čolić, & Marija Stanković. This manuscript was first posted as a preprint on bioRxiv to facilitate rapid dissemination and receive early feedback prior to formal peer review. The authors would like to thank the reviewers for their constructive criticism and valuable suggestions.

