## Supplement for "Negative affective traits moderate tDCS effects on memory"

**Supplementary material**

**Supplement 1.** E-filed models

The estimation of induced e-fields was performed using SimNIBS v4.5 (open-source software package that models current flow and induced e-fields based on finite element methods) on the standardized average head model (derived from high-resolution structural MRI scans). Default tissue conductivity values were applied, while electrode shape, size, and stimulation intensity were specified for each experiment and montage according to the experimental protocols. For each montage, the estimated electric field was extracted at predefined regions of interest (ROIs) based on MNI coordinates: PPC (±47, −71, 34) and DLPFC (±42, 44, 38). Figures below illustrate each of the e-filed model estimates. Across experiments, the estimated e-field values for respective ROIs were as follows:

- Experiment 1: left PPC = 0.267 V/m
- Experiment 2: right PPC = 0.261 V/m
- Experiment 3: left PPC = 0.320 V/m; left DLPFC = 0.245 V/m
- Experiment 4: right PPC = 0.313 V/m; right DLPFC = 0.239 V/m
- Experiment 5: left PPC = 0.267 V/m; left DLPFC = 0.204 V/m
- Experiment 6: left PPC = 0.274 V/m

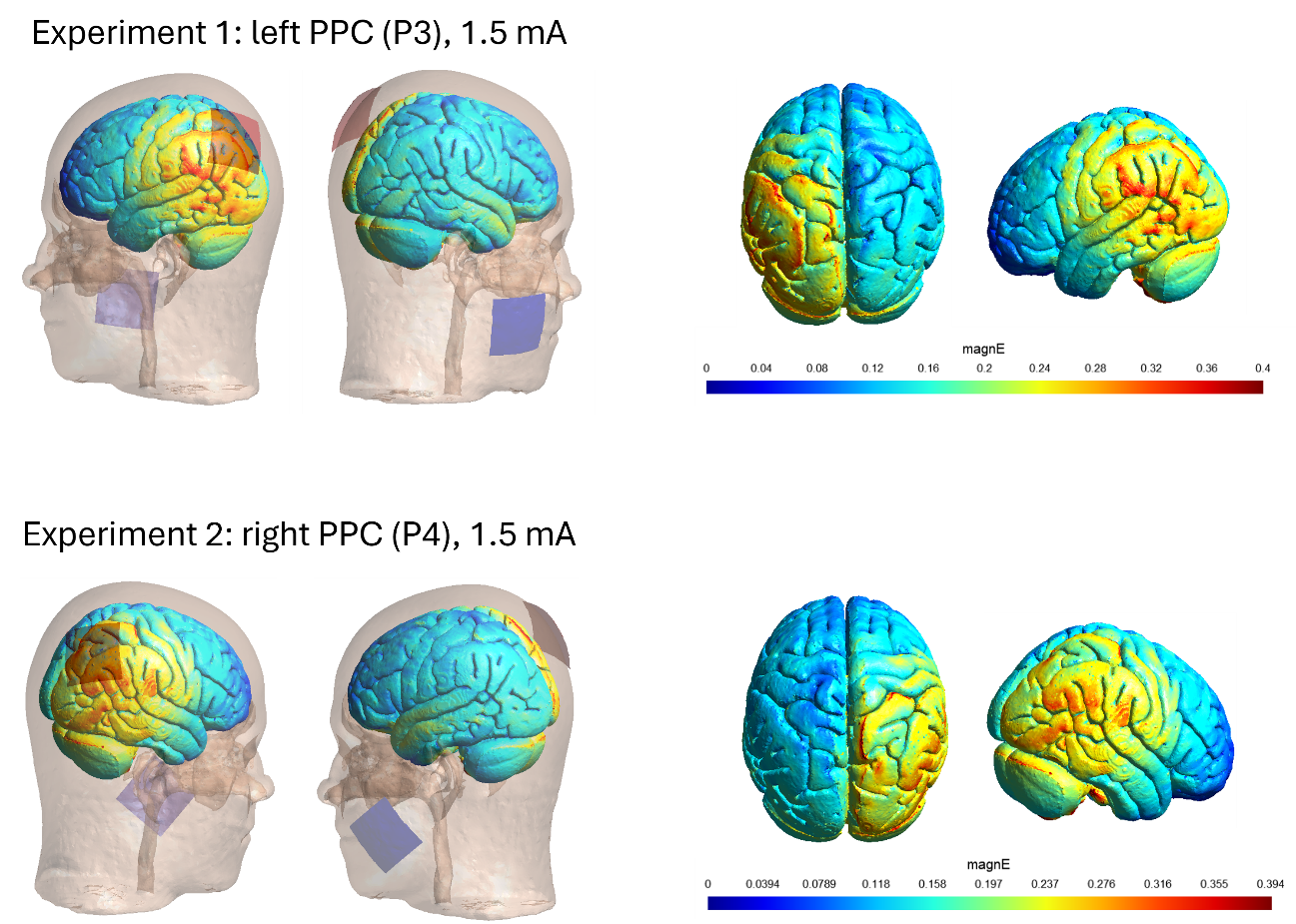

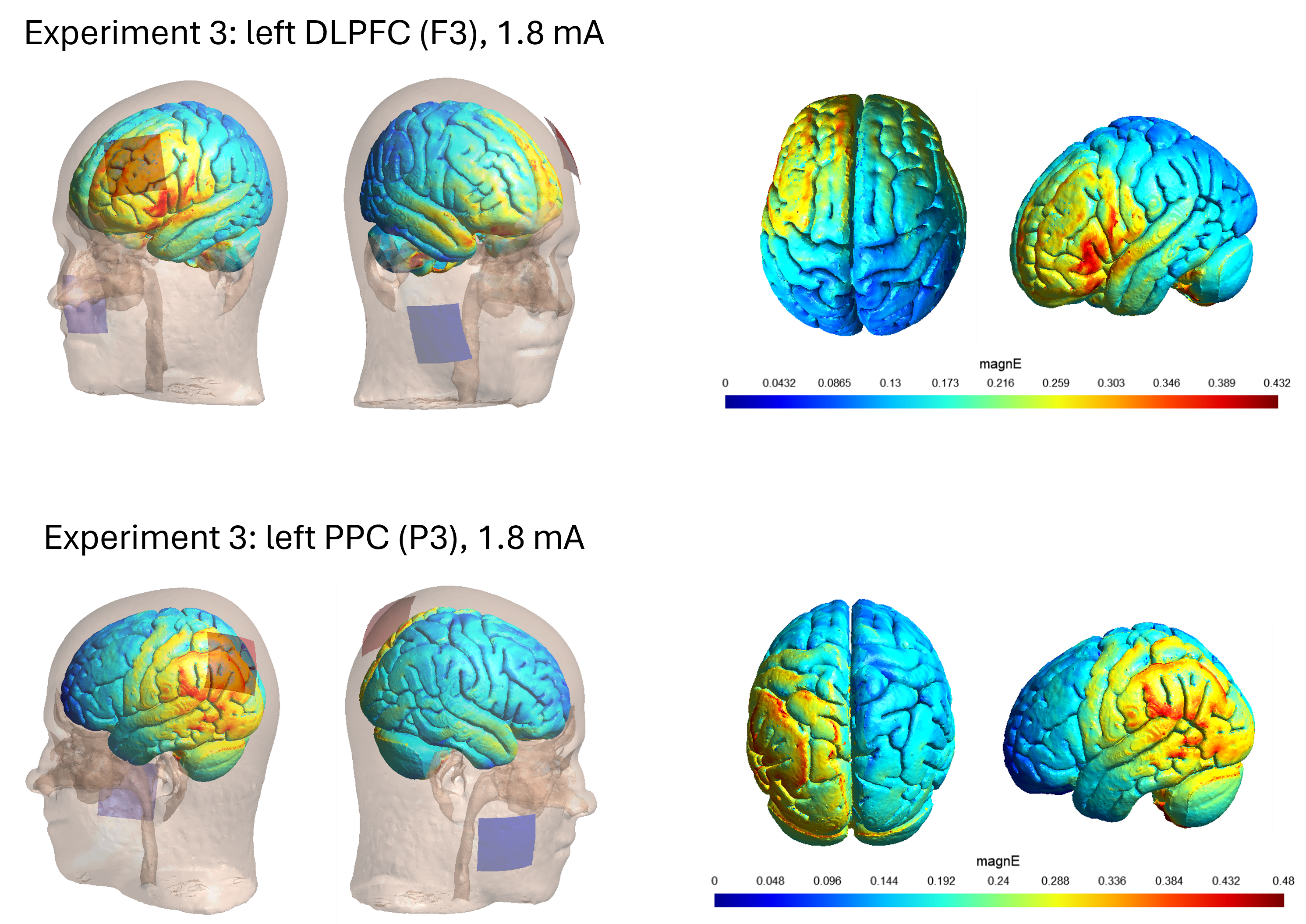

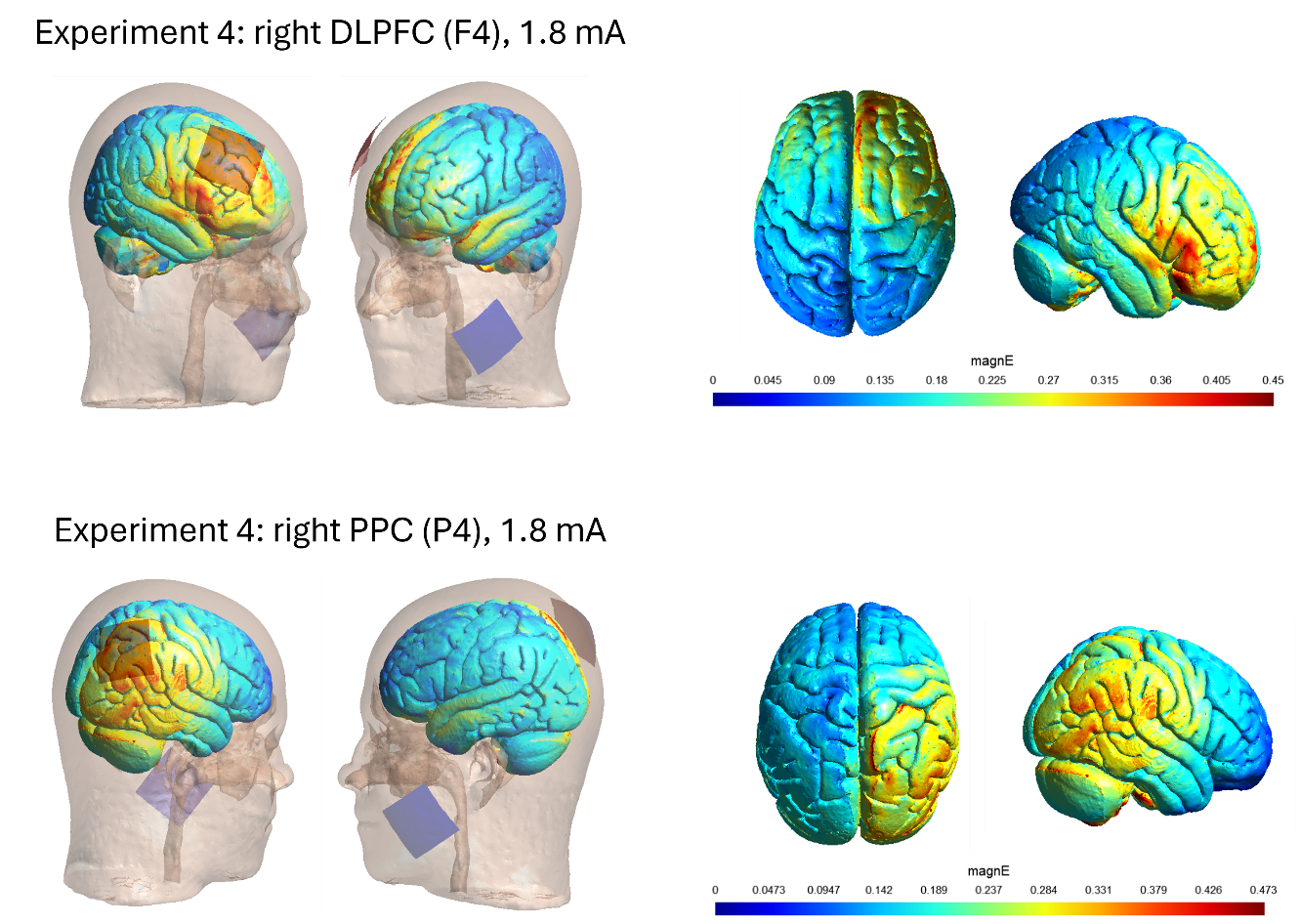

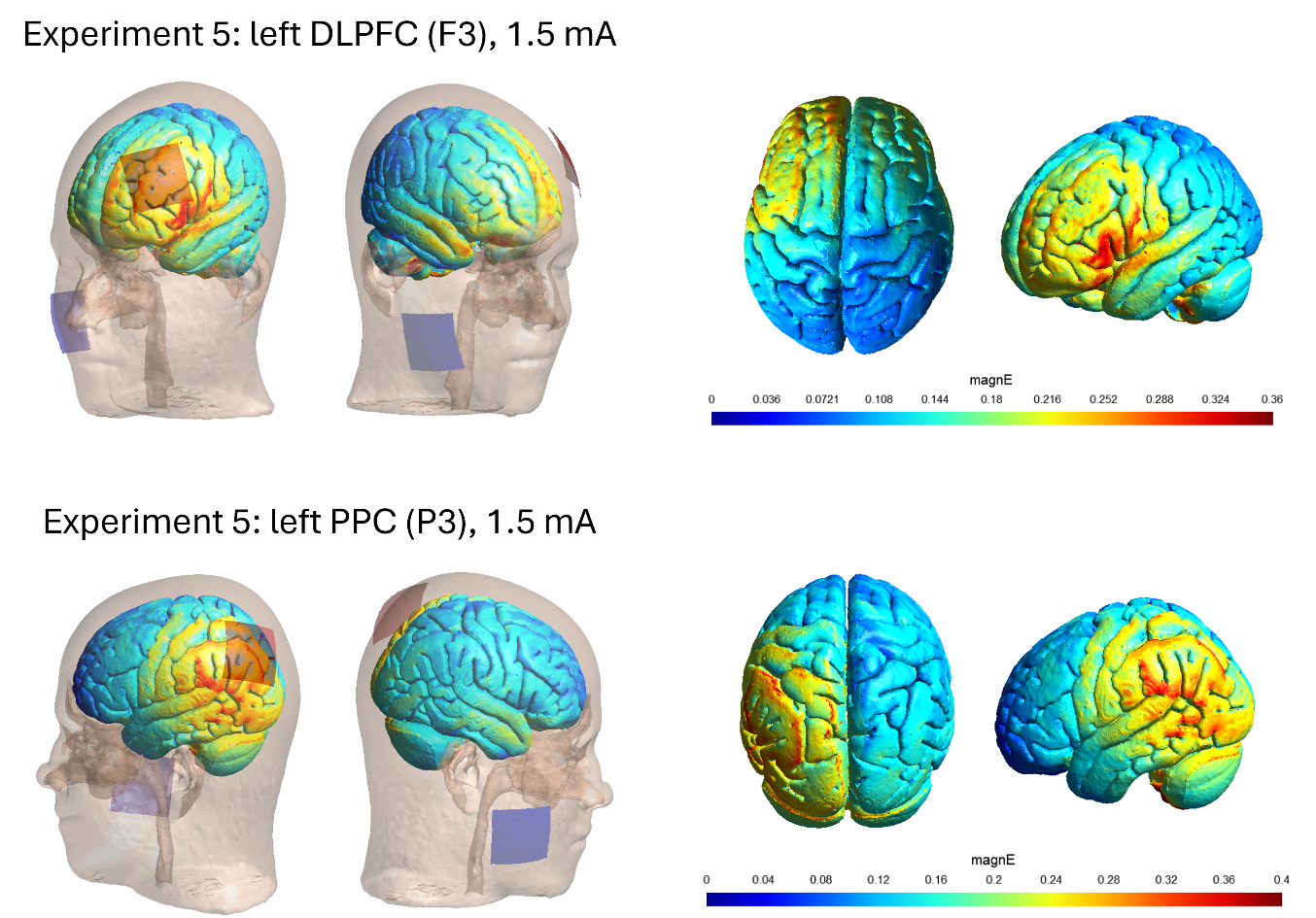

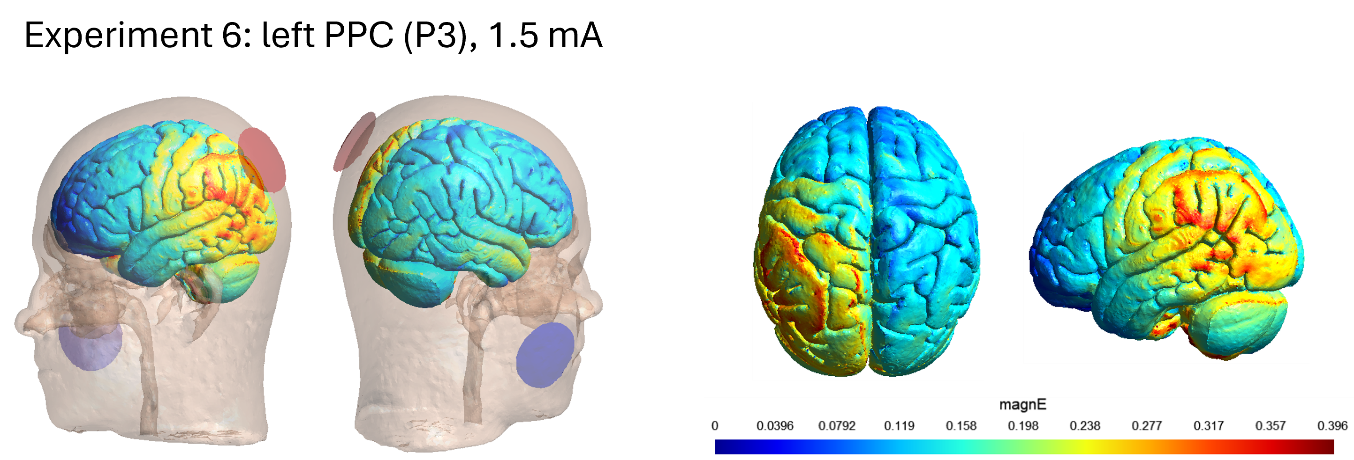

**Supplement 2.** DASS-21 descriptive statistics and distributions

Table S2.1.

*Descriptive statistics for DASS-21 subscales and overall score*

| *Cognitive domain* |  | *Affective domain* | *Sham*  *(M±SD)* | *PPC*  *(M±SD)* | *DLPFC*  *(M±SD)* | *Trait-level*  *(M±SD)* |
| --- | --- | --- | --- | --- | --- | --- |
| WM | *Session-level* | depression | 0.249±0.402 | 0.224±0.425 | 0.218±0.325 | 0.231±0.347 |
|  |  | anxiety | 0.227±0.351 | 0.227±0.348 | 0.268±0.441 | 0.240±0.301 |
|  |  | stress | 0.734±0.619 | 0.617±0.559 | 0.669±0.615 | 0.673±0.520 |
|  |  | DASS overall | 0.403±0.389 | 0.356±0.363 | 0.385±0.388 | 0.381±0.338 |
|  | *State-level* | depression | 0.019±0.192 | -0.006±0.158 | -0.013±0.160 | / |
|  |  | anxiety | -0.014±0.224 | -0.014±0.221 | 0.027±0.258 | / |
|  |  | stress | 0.060±0.341 | -0.056±0.257 | -0.004±0.280 | / |
|  |  | DASS overall | 0.022±0.194 | -0.025±0.141 | 0.003±0.182 | / |
| AM | *Session-level* | depression | 0.317±0.443 | 0.414±0.561 | / | 0.366±0.459 |
|  |  | anxiety | 0.291±0.422 | 0.291±0.441 | / | 0.291±0.389 |
|  |  | stress | 0.772±0.660 | 0.769±0.646 | / | 0.771±0.594 |
|  |  | DASS overall | 0.460±0.426 | 0.492±0.476 | / | 0.476±0.413 |
|  | *State-level* | depression | -.049±.212 | .049±.212 | / | / |
|  |  | anxiety | .000±.188 | .000±.188 | / | / |
|  |  | stress | .002±.271 | -.002±.271 | / | / |
|  |  | DASS overall | -.016±.185 | .016±.185 | / | / |

*Figure S2.1.* Distributions of state- and trait-level depression, anxiety, stress, and overall DASS score for WM data set

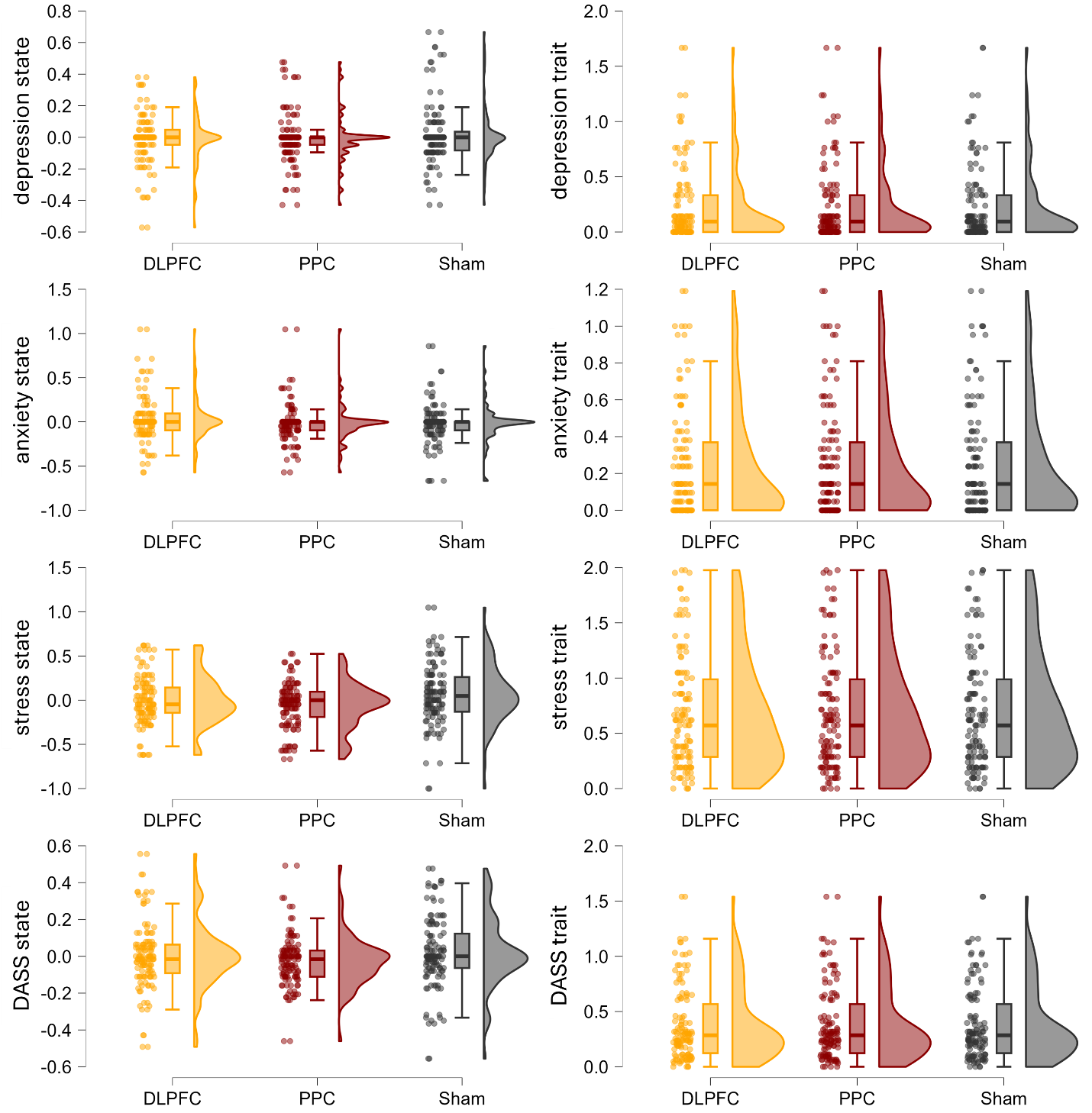

*Figure S2.2.* Distributions of state- and trait-level depression, anxiety, stress, and overall DASS score for AM data set

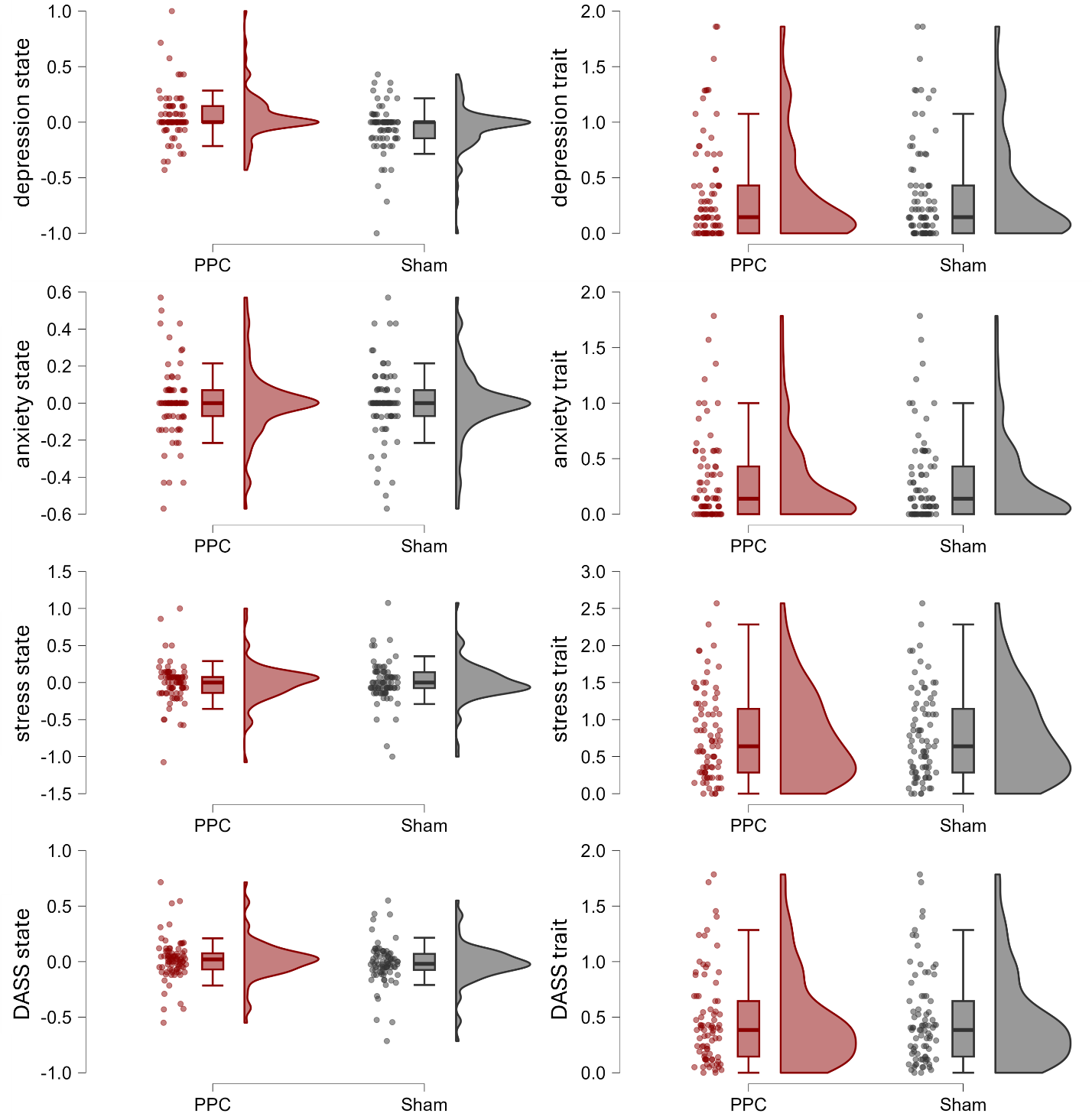

**Supplement 3.** Descriptive statistics for WM and AM (reproduced from the original publications)

Table S3.1.

*Descriptive statistics for WM and AM by condition and task type*

| *Cognitive domain* | *Task* | *Sham*  *(M±SD)* | *PPC*  *(M±SD)* | *DLPFC*  *(M±SD)* |
| --- | --- | --- | --- | --- |
| WM | Verbal 3-back | -0.012±0.157 | 0.011±0.131 | 0.002±0.136 |
|  | Spatial 3-back | -0.020±0.180 | 0.025±0.167 | -0.004±0.180 |
|  | Overall | -0.016±0.156 | 0.018±0.133 | -0.001±0.140 |
| AM | Face-word | 0.300±0.123 | 0.375±0.139 | / |
|  | Object location | 0.457±0.185 | 0.532±0.202 | / |
|  | stAM | 0.663±0.152 | 0.699±0.140 | / |
|  | Overall | 0.520±0.216 | 0.576±0.206 | / |

*Note.* WM – hit rate centered by session order; AM – proportion of successful cued-recall

*Figure S3.1.* Distributions of WM and AM scores

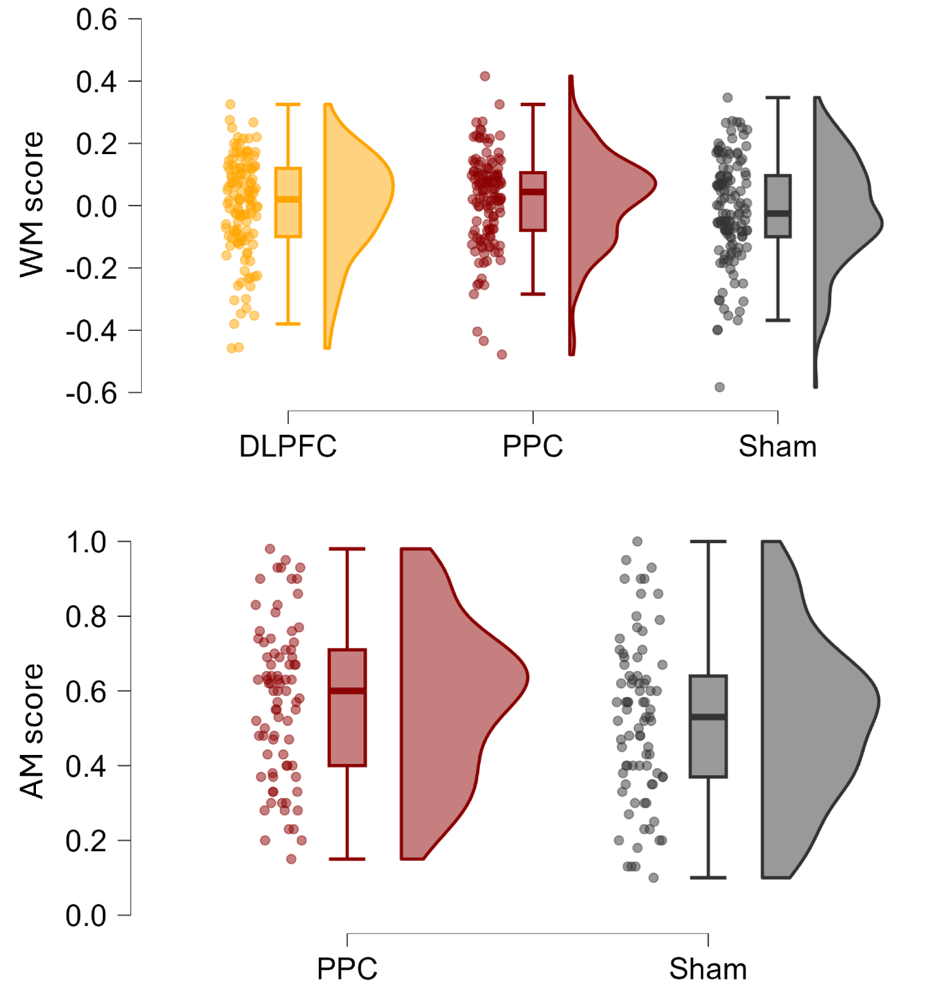

**Supplement 4.** Variance decomposition

**Table S4.1.**

Variance decomposition

| Cognitive  domain | Area  stimulated | Affective  domain | Reliable  variance  (α_max_) | Total variance | | | % within  reliable variance | |
| --- | --- | --- | --- | --- | --- | --- | --- | --- |
|  |  |  |  | Trait-like  (*r_tt_*) | State-like  (α_max_ – *r_tt_*) | Error  (1 – α_max_) | Trait-like  (*r_tt_* _/_ α_max_) | State-like  1 – (*r_tt_* _/_ α_max_) |
| WM | PPC | depression | .879 | .714 | .165 | .121 | 60.2 | 18.8 |
|  |  | anxiety | .759 | .460 | .299 | .241 | 28.3 | 39.4 |
|  |  | stress | .845 | .592 | .253 | .155 | 42.2 | 29.9 |
|  |  | DASS total | .895 | .712 | .183 | .105 | 58.1 | 20.4 |
|  | DLPFC | depression | .847 | .639 | .208 | .153 | 52.5 | 24.6 |
|  |  | anxiety | .861 | .427 | .434 | .139 | 24.4 | 50.4 |
|  |  | stress | .859 | .575 | .284 | .141 | 39.1 | 33.1 |
|  |  | DASS total | .906 | .596 | .310 | .094 | 40.7 | 34.2 |
| AM | PPC | depression | .902 | .667 | .235 | .098 | 52.6 | 26.1 |
|  |  | anxiety | .836 | .622 | .214 | .164 | 48.2 | 25.6 |
|  |  | stress | .888 | .656 | .232 | .112 | 49.9 | 26.1 |
|  |  | DASS total | .929 | .670 | .259 | .071 | 49.2 | 27.9 |

**Supplement 5.** Main effects of tDCS on WM in a model with state-level only moderators

The tDCS effects on WM in models where only state-level negative affectivity is considered as moderator (*WM_score ~ stimulation_condition * state + (1 | ID)*). The main effects of tDCS on WM in the model with state-level **depression** as moderator: *F*_(1,312.12)_ = 5.779, *p* = .017, η_p_^2^ = .02. The main effects of tDCS on WM in the model with state-level **anxiety** as moderator *F*_(1,312.25)_ = 5.580, *p* = .019, η_p_^2^ = .02. The main effects of tDCS on WM in the model with state-level **stress** as moderator *F*_(1,312.95)_ = 5.329, *p* = .022, η_p_^2^ = .02. The main effects of tDCS on WM in the model with state-level **DASS total score** as moderator: *F*_(1,312.47)_ = 5.237, *p* = .023, η_p_^2^ = .02.

**Supplement 6**. Exploratory models with additional factors predicting the effects on WM

Table S6.1.

*The effects of condition (DLPFC, PPC, sham), task, laterality, and negative affectivity on WM*

|  | *Initial model* | | | *Model with state and trait NA* | | |
| --- | --- | --- | --- | --- | --- | --- |
| *Predictors* | *F* | *p* | η_p_^2^ | *F* | *p* | η_p_^2^ |
| condition | 3.998 | .019 | .03 | 0.999 | .370 | .01 |
| task | 0.001 | .972 | .00 | 0.026 | .872 | .00 |
| laterality | 0.001 | .978 | .00 | 0.020 | .887 | .00 |
| depression state | / | / | / | 0.009 | .925 | .00 |
| depression trait | / | / | / | 2.706 | .105 | .04 |
| condition * task | 0.232 | .793 | .00 | 0.030 | .970 | .01 |
| condition * laterality | 4.177 | .016 | .03 | 1.273 | .282 | .00 |
| task * laterality | 0.001 | .978. | .00 | 1.176 | .279 | .01 |
| condition * depression state | / | / | / | 1.695 | .185 | .01 |
| condition * depression trait |  |  |  | 1.801 | .167 | .00 |
| task * depression state | / | / | / | 1.311 | .253 | .00 |
| task * depression trait |  |  |  | 0.569 | .451 | .00 |
| laterality * depression state | / | / | / | 0.081 | .776 | .00 |
| laterality * depression trait |  |  |  | 0.000 | .990 | .00 |
| condition * task * laterality | 0.260 | .771 | .00 | 0.177 | .838 | .00 |
| condition * task * depression state | / | / | / | 0.697 | .499 | .00 |
| condition * task * depression trait |  |  |  | 0.581 | .560 | .00 |
| condition * laterality * depression state | / | / | / | 0.618 | .540 | .00 |
| condition * laterality * depression trait |  |  |  | 1.202 | .302 | .01 |
| task * laterality * depression state | / | / | / | 0.066 | .797 | .00 |
| task * laterality * depression trait |  |  |  | 3.937 | .048 | .01 |
| condition * task * laterality * depression state |  |  |  | 1.217 | .298 | .01 |
| condition * task * laterality * depression trait | / | / | / | 0.019 | .981 | .00 |
| condition | 3.998 | .019 | .03 | 0.010 | .990 | .00 |
| task | 0.001 | .972 | .00 | 0.624 | .430 | .00 |
| laterality | 0.001 | .978 | .00 | 0.148 | .702 | .00 |
| anxiety state | / | / | / | 0.531 | .467 | .00 |
| anxiety trait | / | / | / | 5.395 | .023 | .08 |
| condition * task | 0.232 | .793 | .00 | 0.085 | .919 | .00 |
| condition * laterality | 4.177 | .016 | .03 | 1.827 | .163 | .01 |
| task * laterality | 0.001 | .978. | .00 | 0.001 | .970 | .00 |
| condition * anxiety state | / | / | / | 0.445 | .641 | .00 |
| condition * anxiety trait |  |  |  | 4.990 | .007 | .03 |
| task * anxiety state | / | / | / | 0.297 | .586 | .00 |
| task * anxiety trait |  |  |  | 0.354 | .553 | .00 |
| laterality * anxiety state | / | / | / | 0.067 | .796 | .00 |
| laterality * anxiety trait |  |  |  | 0.604 | .440 | .01 |
| condition * task * laterality | 0.260 | .771 | .00 | 0.669 | .513 | .00 |
| condition * task * anxiety state | / | / | / | 1.664 | .191 | .01 |
| condition * task * anxiety trait |  |  |  | 1.242 | .290 | .01 |
| condition * laterality * anxiety state | / | / | / | 0.698 | .499 | .00 |
| condition * laterality * anxiety trait |  |  |  | 0.732 | .482 | .00 |
| task * laterality * anxiety state | / | / | / | 0.124 | .725 | .00 |
| task * laterality * anxiety trait |  |  |  | 0.310 | .578 | .00 |
| condition * task * laterality * anxiety state |  |  |  | 0.719 | .488 | .01 |
| condition * task * laterality * anxiety trait | / | / | / | 0.407 | .666 | .00 |
| condition | 3.998 | .019 | .03 | 0.477 | .621 | .00 |
| task | 0.001 | .972 | .00 | 1.752 | .187 | .01 |
| laterality | 0.001 | .978 | .00 | 0.009 | .927 | .00 |
| stress state | / | / | / | 0.066 | .798 | .00 |
| stress trait | / | / | / | 5.694 | .020 | .09 |
| condition * task | 0.232 | .793 | .00 | 0.311 | .733 | .00 |
| condition * laterality | 4.177 | .016 | .03 | 1.193 | .305 | .01 |
| task * laterality | 0.001 | .978. | .00 | 0.382 | .537 | .00 |
| condition * stress state | / | / | / | 0.905 | .406 | .01 |
| condition * stress trait |  |  |  | 6.117 | .003 | .04 |
| task * stress state | / | / | / | 0.000 | .994 | .00 |
| task * stress trait |  |  |  | 2.252 | .135 | .01 |
| laterality * stress state | / | / | / | 1.248 | .265 | .00 |
| laterality * stress trait |  |  |  | 0.028 | .868 | .00 |
| condition * task * laterality | 0.260 | .771 | .00 | 0.298 | .742 | .00 |
| condition * task * stress state | / | / | / | 0.381 | .684 | .00 |
| condition * task * stress trait |  |  |  | 1.065 | .346 | .01 |
| condition * laterality * stress state | / | / | / | 0.300 | .741 | .00 |
| condition * laterality * stress trait |  |  |  | 0.426 | .654 | .00 |
| task * laterality * stress state | / | / | / | 0.007 | .931 | .00 |
| task * laterality * stress trait |  |  |  | 0.879 | .349 | .00 |
| condition * task * laterality * stress state |  |  |  | 0.771 | .463 | .01 |
| condition * task * laterality * stress trait | / | / | / | 0.132 | .877 | .01 |
| condition | 3.998 | .019 | .03 | 0.283 | .754 | .00 |
| task | 0.001 | .972 | .00 | 0.936 | .334 | .00 |
| laterality | 0.001 | .978 | .00 | 0.001 | .974 | .00 |
| DASS state | / | / | / | 0.004 | .951 | .00 |
| DASS trait | / | / | / | 6.262 | .015 | .09 |
| condition * task | 0.232 | .793 | .00 | 0.209 | .811 | .00 |
| condition * laterality | 4.177 | .016 | .03 | 0.886 | .413 | .01 |
| task * laterality | 0.001 | .978. | .00 | 0.477 | .491 | .00 |
| condition * DASS state | / | / | / | 1.605 | .202 | .01 |
| condition * DASS trait |  |  |  | 6.087 | .003 | .04 |
| task * DASS state | / | / | / | 0.074 | .785 | .00 |
| task * DASS trait |  |  |  | 0.281 | .597 | .00 |
| laterality * DASS state | / | / | / | 0.519 | .472 | .00 |
| laterality * DASS trait |  |  |  | 0.018 | .894 | .00 |
| condition * task * laterality | 0.260 | .771 | .00 | 0.399 | .672 | .00 |
| condition * task * DASS state | / | / | / | 0.821 | .441 | .01 |
| condition * task * DASS trait |  |  |  | 1.475 | .231 | .01 |
| condition * laterality * DASS state | / | / | / | 0.590 | .555 | .00 |
| condition * laterality * DASS trait |  |  |  | 0.869 | .421 | .01 |
| task * laterality * DASS state | / | / | / | 0.415 | .520 | .00 |
| task * laterality * DASS trait |  |  |  | 2.019 | .156 | .01 |
| condition * task * laterality * DASS state |  |  |  | 2.175 | .116 | .02 |
| condition * task * laterality * DASS trait | / | / | / | 0.254 | .776 | .00 |

Table S6.2.

*The effects of condition (DLPFC, PPC, sham), task, protocol (offline vs online), and negative affectivity on WM*

|  | *Initial model* | | | *Model with state and trait NA* | | |
| --- | --- | --- | --- | --- | --- | --- |
| *Predictors* | *F* | *p* | η_p_^2^ | *F* | *p* | η_p_^2^ |
| condition | 2.385 | .094 | .02 | 0.403 | .669 | .00 |
| task | 0.001 | .982 | .00 | 0.142 | .707 | .00 |
| protocol | 0.000 | .991 | .00 | 0.067 | .797 | .00 |
| depression state | / | / | / | 0.435 | .510 | .00 |
| depression trait | / | / | / | 4.520 | .038 | .07 |
| condition * task | 0.584 | .558 | .00 | 0.132 | .877 | .00 |
| condition * protocol | 1.500 | .225 | .01 | 1.261 | .285 | .01 |
| task * protocol | 0.000 | .993 | .00 | 0.736 | .392 | .00 |
| condition * depression state | / | / | / | 4.156 | .016 | .02 |
| condition * depression trait | / | / | / | 1.648 | .194 | .01 |
| task * depression state | / | / | / | 0.650 | .421 | .00 |
| task * depression trait | / | / | / | 0.925 | .337 | .00 |
| protocol * depression state | / | / | / | 1.021 | .313 | .00 |
| protocol * depression trait | / | / | / | 0.382 | .539 | .01 |
| condition * task * protocol | 0.579 | .561 | .00 | 0.425 | .655 | .00 |
| condition * task * depression state | / | / | / | 0.972 | .380 | .01 |
| condition * task * depression trait | / | / | / | 0.477 | .621 | .00 |
| condition * protocol * depression state | / | / | / | 1.474 | .231 | .01 |
| condition * protocol * depression trait | / | / | / | 0.247 | .781 | .00 |
| task * protocol * depression state | / | / | / | 0.039 | .843 | .00 |
| task * protocol * depression trait | / | / | / | 2.195 | .140 | .01 |
| condition * task * protocol * depression state | / | / | / | 1.294 | .276 | .01 |
| condition * task * protocol * depression trait | / | / | / | 2.757 | .065 | .02 |
| condition | 2.385 | .094 | .02 | 0.247 | .782 | .00 |
| task | 0.001 | .982 | .00 | 0.606 | .437 | .00 |
| protocol | 0.000 | .991 | .00 | 0.182 | .671 | .00 |
| anxiety state | / | / | / | 0.065 | .799 | .00 |
| anxiety trait | / | / | / | 3.769 | .057 | .06 |
| condition * task | 0.584 | .558 | .00 | 0.237 | .789 | .00 |
| condition * protocol | 1.500 | .225 | .01 | 0.797 | .452 | .01 |
| task * protocol | 0.000 | .993 | .00 | 0.031 | .861 | .00 |
| condition * anxiety state | / | / | / | 2.322 | .100 | .01 |
| condition * anxiety trait | / | / | / | 6.923 | .001 | .05 |
| task * anxiety state | / | / | / | 0.114 | .736 | .00 |
| task * anxiety trait | / | / | / | 0.860 | .355 | .00 |
| protocol * anxiety state | / | / | / | 1.284 | .258 | .00 |
| protocol * anxiety trait | / | / | / | 1.169 | .284 | .02 |
| condition * task * protocol | 0.579 | .561 | .00 | 0.605 | .547 | .00 |
| condition * task * anxiety state | / | / | / | 1.288 | .277 | .01 |
| condition * task * anxiety trait | / | / | / | 1.503 | .224 | .01 |
| condition * protocol * anxiety state | / | / | / | 1.584 | .207 | .00 |
| condition * protocol * anxiety trait | / | / | / | 0.421 | .657 | .00 |
| task * protocol * anxiety state | / | / | / | 1.371 | .243 | .00 |
| task * protocol * anxiety trait | / | / | / | 0.413 | .521 | .00 |
| condition * task * protocol * anxiety state | / | / | / | 0.610 | .544 | .00 |
| condition * task * protocol * anxiety trait | / | / | / | 1.083 | .340 | .01 |
| condition | 2.385 | .094 | .02 | 0.968 | .381 | .01 |
| task | 0.001 | .982 | .00 | 2.417 | .121 | .01 |
| protocol | 0.000 | .991 | .00 | 0.112 | .740 | .00 |
| stress state | / | / | / | 0.070 | .791 | .00 |
| stress trait | / | / | / | 6.334 | .015 | .10 |
| condition * task | 0.584 | .558 | .00 | 0.909 | .404 | .01 |
| condition * protocol | 1.500 | .225 | .01 | 0.124 | .883 | .00 |
| task * protocol | 0.000 | .993 | .00 | 0.566 | .453 | .00 |
| condition * stress state | / | / | / | 1.045 | .353 | .01 |
| condition * stress trait | / | / | / | 4.425 | .013 | .03 |
| task * stress state | / | / | / | 0.001 | .981 | .00 |
| task * stress trait | / | / | / | 3.941 | .048 | .01 |
| protocol * stress state | / | / | / | 0.959 | .328 | .00 |
| protocol * stress trait | / | / | / | 0.092 | .762 | .00 |
| condition * task * protocol | 0.579 | .561 | .00 | 0.808 | .447 | .01 |
| condition * task * stress state | / | / | / | 0.270 | .764 | .00 |
| condition * task * stress trait | / | / | / | 1.859 | .158 | .01 |
| condition * protocol * stress state | / | / | / | 0.796 | .452 | .00 |
| condition * protocol * stress trait | / | / | / | 0.110 | .896 | .00 |
| task * protocol * stress state | / | / | / | 1.215 | .271 | .00 |
| task * protocol * stress trait | / | / | / | 0.435 | .510 | .00 |
| condition * task * protocol * stress state | / | / | / | 2.189 | .114 | .02 |
| condition * task * protocol * stress trait | / | / | / | 0.992 | .372 | .01 |
| condition | 2.385 | .094 | .02 | 0.584 | .558 | .00 |
| task | 0.001 | .982 | .00 | 1.493 | .223 | .01 |
| protocol | 0.000 | .991 | .00 | 0.204 | .653 | .00 |
| DASS state | / | / | / | 0.086 | .769 | .00 |
| DASS trait | / | / | / | 6.893 | .011 | .10 |
| condition * task | 0.584 | .558 | .00 | 0.578 | .562 | .00 |
| condition * protocol | 1.500 | .225 | .01 | 0.314 | .731 | .00 |
| task * protocol | 0.000 | .993 | .00 | 0.558 | .456 | .00 |
| condition * DASS state | / | / | / | 3.087 | .047 | .02 |
| condition * DASS trait | / | / | / | 5.453 | .005 | .04 |
| task * DASS state | / | / | / | 0.660 | .417 | .00 |
| task * DASS trait | / | / | / | 2.658 | .104 | .01 |
| protocol * DASS state | / | / | / | 0.049 | .826 | .00 |
| protocol * DASS trait | / | / | / | 0.552 | .460 | .01 |
| condition * task * protocol | 0.579 | .561 | .00 | 0.869 | .420 | .01 |
| condition * task * DASS state | / | / | / | 0.334 | .717 | .00 |
| condition * task * DASS trait | / | / | / | 1.320 | .269 | .01 |
| condition * protocol * DASS state | / | / | / | 2.012 | .135 | .01 |
| condition * protocol * DASS trait | / | / | / | 0.104 | .902 | .00 |
| task * protocol * DASS state | / | / | / | 0.337 | .562 | .00 |
| task * protocol * DASS trait | / | / | / | 0.837 | .361 | .00 |
| condition * task * protocol * DASS state | / | / | / | 2.012 | .136 | .01 |
| condition * task * protocol * DASS trait | / | / | / | 1.353 | .260 | .01 |

**Supplement 7**. Exploratory models with additional factors predicting the effects on AM

Table S7.1.

*The effects of condition (PPC vs sham), task, and negative affectivity on* ***AM***

|  | *Initial model* | | | *Model with state and trait NA* | | |
| --- | --- | --- | --- | --- | --- | --- |
| *Predictors* | *F* | *p* | η_p_^2^ | *F* | *p* | η_p_^2^ |
| condition | 17.756 | <.001 | .19 | 18.458 | <.001 | .20 |
| task | 39.925 | <.001 | .51 | 25.104 | <.001 | .41 |
| depression state | / | / | / | 1.727 | .193 | .02 |
| depression trait | / | / | / | 0.215 | .644 | .00 |
| condition * task | 0.981 | .380 | 02 | 3.746 | .028 | .09 |
| condition * depression state | / | / | / | 0.092 | .763 | .00 |
| condition * depression trait | / | / | / | 2.641 | .109 | .04 |
| task * depression state | / | / | / | 0.771 | .466 | .02 |
| task * depression trait | / | / | / | 0.102 | .903 | .00 |
| condition * task * depression state | / | / | / | 0.238 | .789 | .01 |
| condition * task * depression trait | / | / | / | 3.083 | .052 | .08 |
| condition | 17.756 | <.001 | .19 | 13.043 | <.001 | .15 |
| task | 39.925 | <.001 | .51 | 26.567 | <.001 | .42 |
| anxiety state | / | / | / | 0.167 | .684 | .00 |
| anxiety trait | / | / | / | 2.577 | .113 | .03 |
| condition * task | 0.981 | .380 | 02 | 2.154 | .123 | .06 |
| condition * anxiety state | / | / | / | 1.261 | .265 | .02 |
| condition * anxiety trait | / | / | / | 0.716 | .400 | .01 |
| task * anxiety state | / | / | / | 0.664 | .518 | .02 |
| task * anxiety trait | / | / | / | 0.318 | .729 | .01 |
| condition * task * anxiety state | / | / | / | 1.100 | .338 | .03 |
| condition * task * anxiety trait | / | / | / | 2.632 | .079 | .07 |
| condition | 17.756 | <.001 | .19 | 8.331 | .005 | .10 |
| task | 39.925 | <.001 | .51 | 16.529 | <.001 | .31 |
| stress state | / | / | / | 0.484 | .489 | .01 |
| stress trait | / | / | / | 0.064 | .800 | .00 |
| condition * task | 0.981 | .380 | 02 | 0.681 | .509 | .02 |
| condition * stress state | / | / | / | 0.125 | .725 | .00 |
| condition * stress trait | / | / | / | 0.321 | .573 | .00 |
| task * stress state | / | / | / | 1.554 | .218 | .04 |
| task * stress trait | / | / | / | 0.471 | .626 | .01 |
| condition * task * stress state | / | / | / | 1.081 | .345 | .03 |
| condition * task * stress trait | / | / | / | 0.298 | .744 | .01 |
| condition | 17.756 | <.001 | .19 | 10.972 | .001 | .13 |
| task | 39.925 | <.001 | .51 | 18.798 | <.001 | .34 |
| DASS state | / | / | / | 0.537 | .466 | .01 |
| DASS trait | / | / | / | 0.562 | .456 | .01 |
| condition * task | 0.981 | .380 | 02 | 2.198 | .118 | .06 |
| condition * DASS state | / | / | / | 0.065 | .799 | .00 |
| condition * DASS trait | / | / | / | 0.530 | .469 | .01 |
| task * DASS state | / | / | / | 0.720 | .490 | .02 |
| task * DASS trait | / | / | / | 0.263 | .769 | .01 |
| condition * task * DASS state | / | / | / | 0.613 | .544 | .02 |
| condition * task * DASS trait | / | / | / | 1.593 | .210 | .04 |

Table S7.2.

*The effects of condition (PPC vs sham), laterality, and negative affectivity on* ***AM***

|  | *Initial model* | | | *Model with state and trait NA* | | |
| --- | --- | --- | --- | --- | --- | --- |
| *Predictors* | *F* | *p* | η_p_^2^ | *F* | *p* | η_p_^2^ |
| condition | 14.806 | <.001 | .16 | 18.525 | <.001 | .20 |
| laterality | 2.092 | .152 | .03 | 0.717 | .400 | .01 |
| depression state | / | / | / | 2.615 | .110 | .03 |
| depression trait | / | / | / | 0.063 | .802 | 00 |
| condition * laterality | 0.594 | .443 | .01 | 6.657 | .012 | .08 |
| condition * depression state | / | / | / | 0.000 | .995 | .00 |
| condition * depression trait | / | / | / | 4.982 | .029 | .06 |
| laterality * depression state | / | / | / | 1.506 | .224 | .02 |
| laterality * depression trait | / | / | / | 0.003 | .954 | .00 |
| condition * laterality * depression state | / | / | / | 0.128 | .721 | .00 |
| condition * laterality * depression trait | / | / | / | 6.182 | .015 | .08 |
| condition | 14.806 | <.001 | .16 | 14.055 | <.001 | .16 |
| laterality | 2.092 | .152 | .03 | 0.932 | .337 | .01 |
| anxiety state | / | / | / | 0.914 | .342 | .01 |
| anxiety trait | / | / | / | 1.406 | .239 | .02 |
| condition * laterality | 0.594 | .443 | .01 | 3.949 | .051 | .05 |
| condition * anxiety state | / | / | / | 0.309 | .580 | .00 |
| condition * anxiety trait | / | / | / | 2.222 | .140 | .03 |
| laterality * anxiety state | / | / | / | 0.931 | .338 | .01 |
| laterality * anxiety trait | / | / | / | 0.276 | .601 | .00 |
| condition * laterality * anxiety state | / | / | / | 2.220 | .140 | .03 |
| condition * laterality * anxiety trait | / | / | / | 5.424 | .023 | .07 |
| condition | 14.806 | <.001 | .16 | 8.843 | .004 | .11 |
| laterality | 2.092 | .152 | .03 | 2.157 | .146 | .03 |
| stress state | / | / | / | 1.018 | .316 | .01 |
| stress trait | / | / | / | 0.416 | .521 | .01 |
| condition * laterality | 0.594 | .443 | .01 | 1.478 | .228 | .02 |
| condition * stress state | / | / | / | 0.272 | .603 | .00 |
| condition * stress trait | / | / | / | 0.525 | .471 | .01 |
| laterality * stress state | / | / | / | 3.562 | .063 | .05 |
| laterality * stress trait | / | / | / | 0.864 | .356 | .01 |
| condition * laterality * stress state | / | / | / | 0.049 | .826 | .00 |
| condition * laterality * stress trait | / | / | / | 0.601 | .441 | .01 |
| condition | 14.806 | <.001 | .16 | 12.418 | <.001 | .14 |
| laterality | 2.092 | .152 | .03 | 1.177 | .281 | .02 |
| DASS state | / | / | / | 0.732 | .395 | .01 |
| DASS trait | / | / | / | 0.489 | .487 | .01 |
| condition * laterality | 0.594 | .443 | .01 | 4.420 | .039 | .06 |
| condition * DASS state | / | / | / | 0.035 | .851 | .00 |
| condition * DASS trait | / | / | / | 1.524 | .221 | .02 |
| laterality * DASS state | / | / | / | 1.628 | .206 | .02 |
| laterality * DASS trait | / | / | / | 0.169 | .682 | .00 |
| condition * laterality * DASS state | / | / | / | 0.288 | .593 | .00 |
| condition * laterality * DASS trait | / | / | / | 3.480 | .066 | .04 |

*Note.* The statistically significant interaction effects observed in models including the laterality factor may reflect a spurious finding, potentially attributable to the unequal distribution of observations between left- and right-hemisphere stimulation conditions.

Table S7.3.

*The effects of condition (PPC vs sham), protocol (offline vs online), and negative affectivity on* ***AM***

|  | *Initial model* | | | *Model with state and trait NA* | | |
| --- | --- | --- | --- | --- | --- | --- |
| *Predictors* | *F* | *p* | η_p_^2^ | *F* | *p* | η_p_^2^ |
| condition | 15.870 | <.001 | .17 | 12.476 | <.001 | .14 |
| protocol | 59.715 | <.001 | .43 | 37.463 | <.001 | .33 |
| depression state | / | / | / | 2.994 | .088 | .04 |
| depression trait | / | / | / | 0.187 | .667 | .00 |
| condition * protocol | 1.983 | .163 | .02 | 3.633 | .060 | .05 |
| condition * depression state | / | / | / | 0.469 | .495 | .01 |
| condition * depression trait | / | / | / | 1.031 | .313 | .01 |
| protocol * depression state | / | / | / | 1.990 | .162 | .03 |
| protocol * depression trait | / | / | / | 0.374 | .543 | .00 |
| condition * protocol * depression state | / | / | / | 0.385 | .537 | .01 |
| condition * protocol * depression trait | / | / | / | 2.023 | .159 | .03 |
| condition | 15.870 | <.001 | .17 | 9.083 | .004 | .11 |
| protocol | 59.715 | <.001 | .43 | 38.502 | <.001 | .34 |
| anxiety state | / | / | / | 0.034 | .854 | .00 |
| anxiety trait | / | / | / | 2.144 | .147 | .03 |
| condition * protocol | 1.983 | .163 | .02 | 2.236 | .139 | .03 |
| condition * anxiety state | / | / | / | 0.288 | .593 | .00 |
| condition * anxiety trait | / | / | / | 0.001 | .972 | .00 |
| protocol * anxiety state | / | / | / | 0.297 | .587 | .00 |
| protocol * anxiety trait | / | / | / | 0.013 | .909 | .00 |
| condition * protocol * anxiety state | / | / | / | 1.340 | .251 | .02 |
| condition * protocol * anxiety trait | / | / | / | 0.298 | .587 | .00 |
| condition | 15.870 | <.001 | .17 | 9.808 | .002 | .12 |
| protocol | 59.715 | <.001 | .43 | 27.806 | <.001 | .27 |
| stress state | / | / | / | 0.029 | .866 | .00 |
| stress trait | / | / | / | 0.207 | .651 | .00 |
| condition * protocol | 1.983 | .163 | .02 | 1.223 | .272 | .02 |
| condition * stress state | / | / | / | 0.365 | .548 | .00 |
| condition * stress trait | / | / | / | 0.802 | .373 | .01 |
| protocol * stress state | / | / | / | 0.042 | .839 | .00 |
| protocol * stress trait | / | / | / | 1.043 | .310 | .01 |
| condition * protocol * stress state | / | / | / | 1.187 | .279 | .02 |
| condition * protocol * stress trait | / | / | / | 0.045 | .833 | .00 |
| condition | 15.870 | <.001 | .17 | 9.149 | .003 | .11 |
| protocol | 59.715 | <.001 | .43 | 29.390 | <.001 | .28 |
| DASS state | / | / | / | 0.187 | .667 | .00 |
| DASS trait | / | / | / | 0.732 | .395 | .01 |
| condition * protocol | 1.983 | .163 | .02 | 2.296 | .134 | .03 |
| condition * DASS state | / | / | / | 0.002 | .965 | .00 |
| condition * DASS trait | / | / | / | 0.351 | .555 | .00 |
| protocol * DASS state | / | / | / | 0.017 | .896 | .00 |
| protocol * DASS trait | / | / | / | 0.637 | .427 | .01 |
| condition * protocol * DASS state | / | / | / | 1.143 | .288 | .02 |
| condition * protocol * DASS trait | / | / | / | 0.437 | .511 | .01 |
